# Comparative genomic analyses of trans-ithmanian reef fishes reveals different molecular targets of environmental adaptation in different families

**DOI:** 10.1101/2025.10.28.684510

**Authors:** Eirlys E. Tysall, Carlos F. Arias, Marc P. Hoeppner, Paul B. Frandsen, Claire B. Tracy, Moisés A. Bernal, Oscar Puebla, W. Owen McMillan, Andrea Manica

## Abstract

The extent to which selection acts upon the same molecular targets when faced with the same environmental changes has important implications for our understanding of the repeatability and predictability of evolutionary processes. Replicated natural experiments are well positioned to provide insight, however often these involve closely related species or populations of the same species. Given that the similarity of genetic background makes the same molecular change more likely, it is essential to evaluate the extent to which molecular parallelism occurs in more distantly related species. The rise of the Isthmus of Panama presents an ideal natural experiment to investigate the prevalence of parallel molecular changes in distantly related lineages exposed to the same environmental regimes of temperature, salinity and primary productivity in the two newly formed oceans. Here, we generate high-quality reference genomes for two replicate pairs of geminate reef fishes, one in the family Pomacentridae and one in the Serranidae. Comparative analyses of positive selection in protein-coding genes and of gene family evolution revealed multiple signals of potentially adaptive divergence between geminate species in response to environmental regime. Common targets of selection between families included MAPK signalling (*MAP3K10*), solute transport (*SLC46A2*) and reproduction (*ZPL3L*), as well as expansions in immune and MAPK related gene families. Overall, however, we found minimal evidence of parallelism between families either at the gene level or when considering higher functional categories. Our findings suggest that phylogenetic constraints may limit the levels of molecular parallelism in distant lineages even when faced with comparable selective pressures.

**Significance Statement:** Understanding the extent to which species exposed to similar environmental conditions may adapt using similar genetic solutions has important implications for our understanding of evolutionary processes and how we may expect species to adapt in the future. However, systems where multiple, independent lineages are faced with the same shift in conditions are rare. The rise of the Isthmus of Panama provides a unique, natural experiment where multiple marine species have been divided and exposed to radically different environmental conditions. Here, we generate genomes for separated species in two distantly related families of reef fishes and compare genome-wide patterns of divergence. Our analyses identified signals of selection in protein-coding genes and shifts in gene family size consistent with adaptation to the distinct environments, however the specific genes and pathways involved predominantly differed between the families, revealing that, even under shared selective pressures, molecular mechanisms may differ across more distant lineages.

## Introduction

Which traits divergent selection acts upon and the ecological conditions driving such selection are fundamental questions relevant to our understanding of adaptive evolution (Kawecki and Ebert 2004). In particular, the degree to which genetic changes may lead to repeatable or predictable adaptations to similar environmental pressures is essential for evolutionary biology (Christin et al. 2010, Stern 2013, Lachapelle et al. 2015, Storz 2016, Bolnick et al. 2018, Hao et al. 2019). While there are several definitions used across the literature to describe convergent versus parallel evolution (Arendt and Reznick 2008, Pearce 2012, Rosenblum et al. 2014, Stuart 2019, Cerca 2023), here, we use the definitions suggested by Bolnick et al. (2018). The latter suggests convergence and parallelism can be distinguished based on geometric patterns of evolutionary trajectories, where convergence refers to the independent evolution from different starting conditions to more similar genotypes or phenotypes in response to comparable selective pressures, while, parallelism refers to independent evolution from similar starting conditions (Bolnick et al. 2018). Such parallelism may be present at the molecular level or be confined to phenotype.

Evaluating the extent to which adaptive parallelism in response to similar selective pressures extends to the molecular level through changes to the same genes or functional group of genes is challenging, particularly in natural populations. Whilst there are striking examples of parallel adaptive changes at the molecular level even to the same amino acid substitutions across more distant lineages (e.g. Musilova et al. 2021), more often these examples occur in closely related species or populations of the same species (Colosimo et al. 2005, Protas et al. 2006, Baxter et al. 2008, Chan et al. 2010, Brown et al. 2019, Moran et al. 2023), whose similar genetic background and likelihood of residing in comparable environments under similar selective pressures may increase the likelihood of similar molecular solutions (Bridgham 2016, Hao et al. 2019). Divergent genetic backgrounds in more distant lineages are less likely to find the same mutation beneficial (Natarajan et al. 2016, Storz 2016, Bohutínská & Peichel 2024). Additionally, where increasing genetic divergence means reduced shared standing genetic variation between lineages, the number of shared beneficial alleles and opportunities for allele reuse will be reduced, as has been shown in adaptation to alpine environments in *Arabidopsis* (Bohutínská et al. 2021) and altitude in tropical butterflies (Montejo-Kovacevich et al. 2022). The predictability of parallel responses in phylogenetically distant species residing in comparable environments may be more restricted to higher functional levels, though appropriate systems to test these hypotheses are limited in marine systems.

The rise of the Isthmus of Panama initiated the separation of multiple, previously connected populations of marine species and sent them on independent evolutionary trajectories for at least the last 3 million years (Coates et al. 2005, O’Dea et al. 2016). The challenge of reconstructing the process of the formation of the Isthmus of Panama has led to some discourse over the timing of its initiation and final closure, however the current consensus suggests the predominance of evidence supports a ∼3 million years ago date for its final closure (O’Dea et al. 2016). For many reef-associated species, the date of their separation may not correspond to the final formation of the isthmus (Knowlton and Weigt 1998, Lessios 2008). The gradual process of the rise of the isthmus likely means the date when gene flow was no longer possible is likely to be the most relevant and one that will be different for different species and lineages. As exemplified in the snapping shrimp (Alpheus spp.) (Knowlton et al. 1993, Knowlton and Weigt 1998), factors such as ecology and life history traits can be important in timings of separation between different species. Termed ‘geminate species’ (Jordan 1908), the importance of these sister species pairs separated by the isthmus (either upon closure or before) as natural experiments in divergence and adaptation has spurred considerable research which has helped identify pairs across multiple taxa (Knowlton and Weigt 1998, Marko 2002, Lessios 2008, Marko and Moran 2009).

Here we consider two geminate species pairs one from the fish family Pomacentridae (damselfishes): *Azurina atrilobata* (scissortail chromis, Pacific) and *Azurina multilineata* (brown chromis, Atlantic) and from the family Serranidae (groupers and seabasses): *Cephalopholis colonus* (Pacific creolefish) and *Paranthias furcifer* (Atlantic creolefish). The geminate species of these families occupy very similar ecological niches; they are found sympatrically at the same depth in the water column, forage during the same time of day and are planktivores feeding on an overlapping diet (Randall 1967). Whilst this indicates a largely shared selection regime, a range of other factors including demographic history, time since divergence, phylogenetic distance and genetic constraints may restrict parallelism at the molecular level (Rosenblum et al. 2014, Hao et al. 2019). Existing phylogenies (Craig and Hastings 2007, Rabosky et al. 2018, McCord et al. 2021) and estimates of molecular divergence in mitochondrial regions (Lessios 2008) indicate divergence times are likely to be different between the geminates of both families with the damselfish diverging earlier. The Pomacentridae and Serranidae families themselves are distantly related, having diverged more than 100 million years ago - a timescale comparable to that of humans and elephants (Kumar et al. 2017). Additionally, the phylogenetic constraints acting on these families are potentially very different. The pomacentrid geminates belong to the Chrominea subfamily of damselfishes which predominantly specialise in pelagic planktivory, with the most likely ancestor of the lineage being planktivorous (McCord et al. 2021). Contrastingly the serranid geminates have had a recent niche shift from the more typical Serranidae niche of nocturnal predator to diurnal planktivore (Ng et al. 2025). In this respect, the rise of the isthmus provides a rare and powerful comparative experimental system for determining the extent to which adaptive change to the same selective pressures may be achieved through the same molecular mechanisms even in distantly related species.

The isthmus not only separated species, but also resulted in dramatic changes in the physical environment in the ocean basins on either side (D’Croz and O’Dea 2007, Lessios 2008). Since the final closure of the isthmus (∼3 mya) considerable differences in most major environmental axes, including temperature, dissolved oxygen, pH and primary productivity, have been in place in the two oceans (Keigwin 1982, Cannariato and Ravelo 1997, D’Croz and Robertson 1997, D’Croz et al. 1999, Haug et al. 2001, O’Dea et al. 2007, 2012, Jain and Collins 2007, D’Croz and O’Dea 2007). Today the Tropical Eastern Pacific (TEP) is characterised by high variability both spatially and temporally. Seasonal upwellings of cool, nutrient-rich water lead to high primary productivity and extreme drops in temperature (D’Croz and Robertson 1997, D’Croz and O’Dea 2007). Conversely, the waters of the Caribbean are much more stable, experience minimal primary productivity, show greater clarity, higher salinity levels and increased temperatures (D’Croz and Robertson 1997, D’Croz et al. 1999). Whilst these distinct changes are set with the final closure, the long and complex process of the isthmus’ formation is thought to have led to environmental shifts in the period before, important in cases where geminate species have split prior to the final closure. Between 4.25 and 3.45 ma, the Caribbean experienced a sharp decline in upwelling, reduced seasonal temperature variation, increased carbonate production, and a collapse in primary productivity (O’Dea et al. 2007, Jackson and O’Dea 2023). By 4.2 ma the salinity contrast between the two oceans had reached those seen today (Haug et al. 2001). Meanwhile in the TEP there is evidence of the presence of seasonal upwellings similar to those seen today from the mid-Pleistocene (O’Dea et al. 2012) and evidence for high surface-level nutrients in the ocean as far back as 5 ma (Cannariato and Ravelo 1997). The well-understood timing and oceanographic effects in this system provide a unique backdrop to explore adaptive divergence in multiple marine species subject to the same change in environmental conditions.

The environmental shifts experienced by marine fishes isolated by the formation of the isthmus have been historically identified as key variables leading to adaptive divergence. In particular, we predict differences in salinity, temperature and nutrient levels to drive adaptations in genomic regions associated with maintenance of osmoregulation (ATPase pumps, ion cotransporters and channels, osmosensors and protein hormones; Manzon 2002, Shimada et al. 2011, Lamichhaney et al. 2012, Berg et al. 2015, Martinez Barrio et al. 2016, Velotta et al. 2022), homeostasis (ion transport, heat shock proteins, and aquaporins; Fangue et al. 2006, Feidantsis et al. 2012, Chu et al. 2021) and metabolic processes (metabolic enzymes, insulin receptor, lipid metabolism regulators, melanocortin receptors and nutrient transporters; Aspiras et al. 2015, Riddle et al. 2018, Li et al. 2024). Furthermore, we know from comparisons of morphology and life history traits of geminates species that there is evidence for repeated phenotypic adaptation in geminate pairs to the different environmental regimes in the TEP and Caribbean (Lessios 1990, Moran 2004, Aguilar-Medrano 2018); however, knowledge of the genetic basis of adaptation in this system is lacking (but see Tracy et al. 2025).

Here we generate high-quality reference genomes for the damselfish geminate species *A. atrilobata* and *A. multilineata* and grouper geminate species *C. colonus* and *P. furcifer*. In addition, we generate reference genomes for the most closely related outgroups of each geminate pair–*Azurina cyanea* (blue chromis) for the damselfish and *Cephalopholis fulva* (coney grouper) for the groupers, both of which are distributed in the Caribbean. Using this data we address the following questions: 1) How have the genomes of the geminate species diverged since their separation? 2) Do the geminate species show evidence of adaptation linked to the distinct environmental conditions in the Caribbean and TEP? 3) To what extent do the two families exhibit parallel genomic responses to these contrasting environments? Taken together, these analyses provide a foundation to understand how genomes of distantly related groups change in response to the same biogeographic event.

## Results

### Genome Assembly Metrics

Metrics for the six assembled genomes, following contaminant detection using BlobTools, are summarised in Table 1. Overall, the assemblies were close to chromosome-level resolution. Most genomes were assembled into fewer than 100 contigs, with 90% of the genome contained within approximately 23 contigs (Supplementary Figure S1) and an average N50 of ∼36 Mb (Table 1). The *A. atrilobata* genome was an exception; it assembled into 434 contigs, with 90% of the genome contained within 61 contigs and an N50 of 26.2 Mb. Genome completeness, assessed using BUSCO, revealed that all assemblies achieved high completeness (>99%) with low percentages of missing predicted orthologs (Supplementary Figure S2). Synteny analysis demonstrated a high degree of synteny between geminate species pairs and their respective outgroups (Supplementary Figure S3). Notably, synteny was stronger between geminate taxa, while comparisons with outgroup taxa revealed a greater number of rearrangements (Supplementary Figure S3).

**Table 1.** Genome assembly metrics for the damselfish and grouper geminate species pairs and outgroups.

| Species | Total Contigs | Genome Size (Mb) | Longest Contig (Mb) | N50 (Mb) | N90 (Mb) |
| --- | --- | --- | --- | --- | --- |
| *A. atrilobata* | 434 | 817.00 | 35.1 | 26.2 | 86.9 |
| *A. multilineata* | 42 | 789.37 | 38.0 | 34.0 | 27.2 |
| *A. cyanea* | 99 | 859.00 | 42.2 | 35.0 | 27.1 |
| *C. colonus* | 60 | 956.00 | 48.1 | 40.5 | 26.6 |
| *P. furcifer* | 77 | 955.00 | 47.8 | 39.2 | 29.1 |
| *C. fulva* | 59 | 1002.00 | 50.5 | 42.5 | 31.3 |

### Molecular Divergence

The damselfish geminate species exhibited a higher number of genomic rearrangements compared to the grouper geminate species, suggesting distinct, lineage-specific evolutionary dynamics. One possible explanation for these differences is that the damselfish lineage may have begun diversifying prior to the final closure of the Isthmus of Panama, resulting in deeper and more complex patterns of genomic divergence. This interpretation is supported by our analysis of 842 orthologous genes, which revealed contrasting divergence patterns between the two groups. Using Dxy, Ks, and molecular clock-based divergence time estimates, both damselfish and grouper species pairs showed right-skewed distributions of Dxy and Ks, with peaks at low values indicating a high level of gene conservation (Supplementary Figure S4). However, damselfish displayed broader distributions in both metrics, suggesting greater divergence or variability in substitution rates. Correspondingly, divergence times were higher in damselfish, with a mean of 12.16 million years ago (MYA) and a mode of 7.74 MYA, compared to a mean of 5.38 MYA and a mode of 3.29 MYA in groupers. These patterns are consistent with an earlier onset of divergence in damselfish and highlight the influence of historical contingency and lineage-specific processes in shaping patterns of genomic evolution.

### Positive selection but no evidence for parallelism

Through our branch-site selection analyses using ETE-Toolkit (Huerta-Cepas et al. 2016), we found evidence of selection across a range of genes in the damselfish and grouper geminates, with a stronger signature of divergent selection in the former when compared to the latter. Additionally, comparisons between the two families revealed minimal overlap in the individual genes or higher functional gene categories under divergent selection. In total we recovered 4279 single copy orthologs present in all ten damselfish species included in the analyses and 3844 between the ten grouper species. Of these we found 1227 were shared between the damselfish and groupers (based on functional annotation). In the Caribbean species’ (*A. multilineata* and *C. furcifer*), we detected significant signatures of selection in 387 genes in the damselfish geminate and 246 genes in grouper geminate. Of these, only thirteen genes were found positively selected in both families, with broad functions including development, cell cycle regulation, signal transduction, and neural function (Table 2). Of note is a mitogen-activated protein kinase gene (*MAP3K10*), part of the c-Jun N-terminal kinases or stress-activated protein kinases (JNK/SAPK) subfamily. This signalling pathway is likely to play a significant role in adaptive responses to various environmental stressors, including thermal, osmotic, and hypoxic challenges (Cowan and Storey 2003).

**Table 2.** Genes identified as under positive selection in both the Caribbean grouper geminate *Paranthias furcifer* and the Caribbean damselfish geminate *Azurina multilineata* at *p* < 0.05. Genes significant at an 10% FDR threshold (Benjamini-Hochberg adjusted p-value < 0.1) are indicated. Gene names and description reported are given by eggnog-mapper annotation.

| Ortholog ID | Gene Name | Gene Description | Significant FDR 10% |  |
| --- | --- | --- | --- | --- |
|  |  |  | *Damselfish* | *Grouper* |
| OG0003785 | *YTHDC1* | YTH domain containing 1 | Y | Y |
| OG0003818 | - | Zona pellucida<br>sperm-binding protein 3-like | Y | Y |
| OG0003830 | - | Nuclear factor of activated<br>T-cells, cytoplasmic | Y | Y |
| OG0004867 | - | Cingulin-like | Y | Y |
| OG0004869 | *RFX7* | Regulatory factor X, 7 | N | N |
| OG0005180 | *TDGF1* | Teratocarcinoma-derived<br>growth | N | Y |
| OG0005247 | *SEZ6* | 6 homolog | N | Y |
| OG0005413 | - | Si ch211-136a13.1 | Y | Y |
| OG0005481 | *map3k10* | Mitogen-activated protein<br>kinase kinase kinase | Y | N |
| OG0005573 | *RCSD1* | RCSD domain containing 1 | Y | Y |
| OG0006979 | *LMOD3* | Leiomodin 3 (fetal) | Y | Y |
| OG0007130 | *CDK11B* | Cyclin-dependent kinase | Y | Y |
| OG0007507 | *GRB7* | Growth factor<br>receptor-bound protein | Y | Y |

Comparatively, in the TEP species’ (*A. atrilobata* and *C. colonus*) we found 531 genes positively selected in the damselfish geminate compared to only 256 in the grouper geminate. For genes under selection in both families in the TEP we found a similar story as in the Caribbean, with just thirteen genes shared (Table 3). These shared genes again cover broad functions generally representing core cellular processes, however several may function in immune response (*POU2AF1*, *TRIM16* and *SKAP1*). Immune cell adaptor *SKAP1* plays multiple roles in T-cell signalling and adaptive immunity (Dadwal et al. 2021, Wohlleben et al. 2023, 2024) and TRIM protein *TRIM16* has been shown to be involved in innate immune response against viral infection in teleosts (van der Aa et al. 2009, Yu et al. 2016, Cho et al. 2022). The role of *POU2AF1* in teleosts is less well understood (Lennard Richard et al., 2007), though in mammals, it is involved in the B-cell response and the production of antibodies (Yeremenko et al. 2021). Additionally, solute carrier *SLC46A2* has been linked to salinity stress in other teleost species (Escobar-Sierra and Lampert 2024b) and reproduction related gene *Zona pellucida sperm-binding protein 3-like* is involved in egg envelope structure and mediation of sperm binding.

**Table 3.** Genes identified as under positive selection in both the Tropical Eastern Pacific grouper geminate *Cephalopholis colonus* and the Caribbean damselfish geminate *Azurina atrilobata* at *p* < 0.05. Genes significant at a 10% FDR threshold (Benjamini-Hochberg adjusted p-value < 0.1) are indicated. Gene names and description reported are given by eggnog-mapper annotation.

| Ortholog ID | Gene Name | Gene Description | Significant FDR 10% |  |
| --- | --- | --- | --- | --- |
|  |  |  | *Damselfish* | *Grouper* |
| OG0004754 | *TICRR* | TOPBP1-interacting checkpoint and replication regulator | Y | Y |
| OG0005179 | *DCC* | DCC netrin 1 receptor | Y | Y |
| OG0005268 | *POU2AF1* | POU class 2 associating factor 1 | Y | Y |
| OG0005289 | - | Alkaline phosphatase | Y | Y |
| OG0005294 | *pax3* | Paired box | Y | Y |
| OG0005560 | *CADM2* | Cell adhesion molecule | Y | Y |
| OG0006688 | - | Cytochrome c oxidase subunit IV isoform 2 | Y | N |
| OG0006806 | *FANCE* | Fanconi anemia complementation group E | Y | N |
| OG0006948 | *CDH26* | Cadherin 26, tandem duplicate<br>1 | N | **Y** |
| OG0007208 | *TRIM16* | Tripartite motif-containing | **Y** | **Y** |
| OG0007487 | *SKAP1* | Src kinase associated<br>phosphoprotein 1 | **Y** | **Y** |
| OG0007529 | *ITGB4* | integrin | N | **Y** |
| OG0008554 | *SLC46A2* | Solute carrier family 46<br>member 2 | **Y** | **Y** |

When considering the next hierarchical level of potential parallelism, the functional enrichment of these positively selected genes, we again found minimal overlap of GO terms between the grouper and damselfish geminates. Between the Caribbean grouper and damselfish species, just one GO term was found functionally enriched in both: positive regulation of transcription by RNA polymerase II (GO:0045944) (Supplementary Table S1). In the TEP we found four terms in common; integrin-mediated signalling pathway (GO:0007229), chromosome segregation (GO:0007059), positive regulation of viral genome replication (GO:0045070) and localization of cell (GO:0051674) (Supplementary Table S2).

Since we found little evidence of parallel responses in the damselfish and grouper geminates at either the gene or functional enrichment level, we investigated how closely the functional enrichment terms in either the TEP or Caribbean clustered together between the groupers and damselfish based on semantic similarity. In this approach, the SimRel (Schlicker et al. 2006) semantic similarity measure calculated between GO terms was visualised using multidimensional scaling, from which clusters of semantically related terms could be formally identified. Clustering of enriched GO terms revealed overlap of GO terms between the groupers and damselfish within semantic clusters, as well as elucidating enrichment clusters potentially indicative of adaptation to the distinct conditions of the TEP and Caribbean (Figure 1; Supplementary Tables S1-S2).

**Figure 1.**
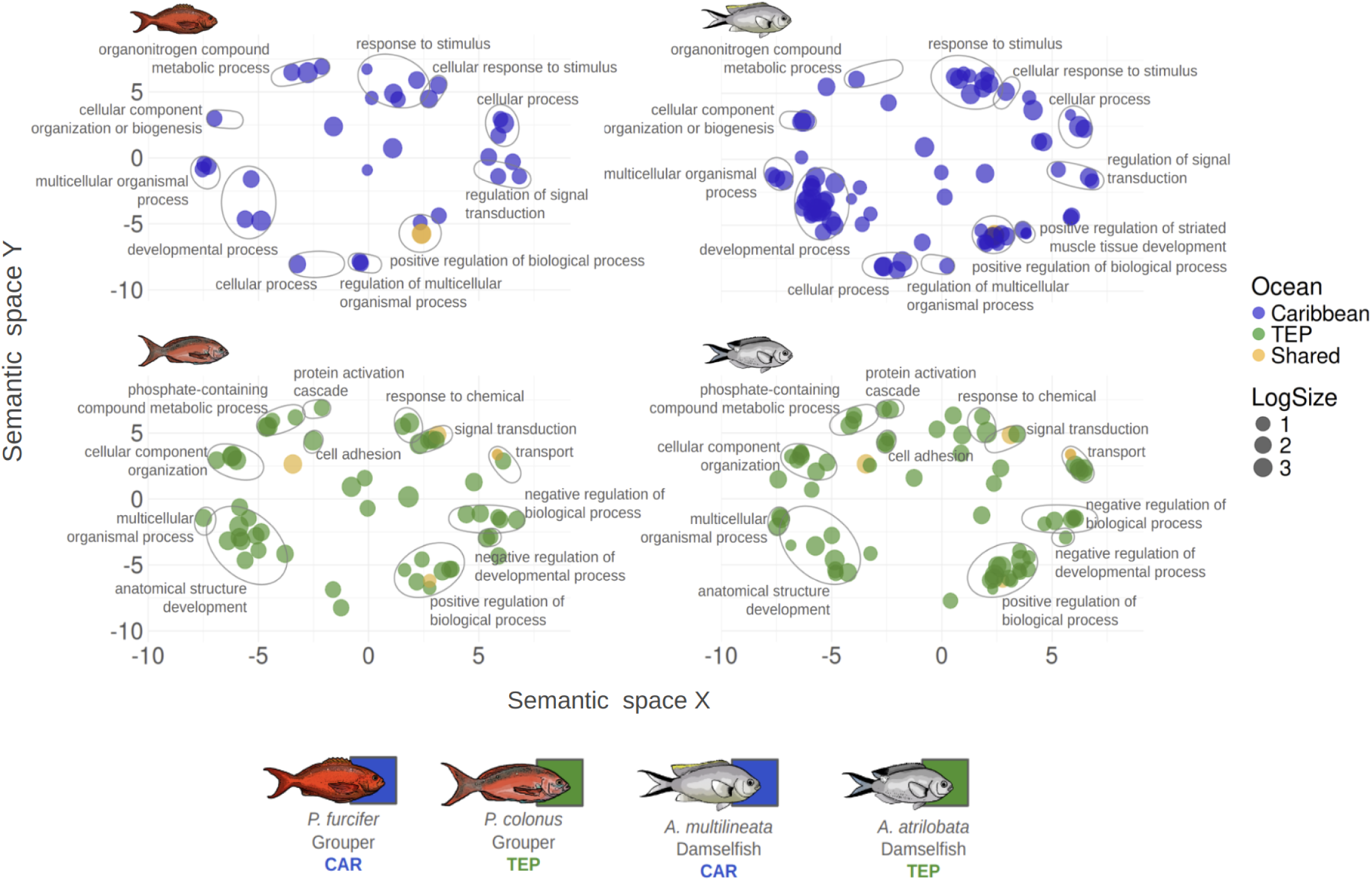
MDS plot of the pairwise semantic similarity of significantly enriched GO terms based on branch-site selection analyses. More semantically similar GO terms will be closer together. Clustering analysis was performed on the two TEP species *A. atrilobata* (damselfish) and *C. colonus* (grouper) and on the two Caribbean species *A. multilineata* (damselfish) and *P. furcifer* (grouper) to identify clusters of similar GO terms between families in the same ocean environment. Clusters are identified with an ellipse and cluster label of the closest common ancestor GO term based on the GO hierarchy. GO terms which are shared between species in the same ocean are marked in orange. LogSize refers to the frequency of the GO term in the underlying EBI Gene Ontology Annotation (GOA) database.

### GO Semantic Clusters in the Caribbean

In the Caribbean, for the clusters which contain GO terms from both families, the ancestor label which best describe these clusters (i.e. the closest common ancestor term of the cluster based on the GO hierarchy) primarily included broad terms indicative of general housekeeping functions such as “Cellular process”, “Developmental process”, “Positive regulation of biological process”, “Multicellular organismal process”, “Regulation of signal transduction”, “Cellular component organization or biogenesis” and “Regulation of multicellular organismal process” (Figure 1; Supplementary Table S1). However, we also see shared clusters in functions which we may expect to be related to adaptation to external environmental conditions: “Response to stimulus”, “Cellular response to stimulus”, “Organonitrogen compound metabolic process”, and “Regulation of signal transduction”. In the groupers, the stimulus response clusters included terms Cellular Response to heat (GO:0034605), Cellular response to calcium ion (GO:0071277), Inflammatory response (GO:0006954), and Cellular response to abiotic stimulus (GO:0071214). In the damselfish we see Cellular response to nutrient levels (GO:0031669), Defence response (GO:0006952), Response to antibiotic (GO:0046677) and Cellular response to xenobiotic stimulus (GO:0071466). This overlapping cluster highlights biological processes that may enable species to cope with various environmental pressures and suggests that both the groupers and damselfish have adapted to the Caribbean environment through responses to various external stimuli, stressors, and nutrient fluctuations which are present.

The shared cluster “Regulation of signal transduction” contains functionally enriched terms involved in signalling pathways and stress responses that may be modulated by external stimuli, especially environmental stressors or chemical signals encountered in the Caribbean. In the groupers this contained Regulation of cAMP-mediated signaling (GO:0043949), Negative regulation of G protein-coupled receptor signaling pathway (GO:0045744) and in the damselfish Regulation of p38MAPK cascade (GO:1900744). Interestingly the p38MAPK cascade is another MAPK subfamily thought to be activated by stress stimuli, specifically playing a role in teleosts in response to osmotic and thermal stress (Kültz and Avila 2001, Feidantsis et al. 2012). Additionally, we also find unclustered but relevant GO term Positive regulation of MAP kinase activity (GO:0043406) significantly enriched in the groupers. This signal, coupled with the shared positive selection on MAPK gene (*MAP3K10*), indicates that this stress induced signalling pathway is a common target of selection in both the groupers and damselfish and, based on its known function, it may be highly likely to play a role in adaptation to environmental conditions in these species.

Finally, in the grouper geminate, we found a cluster of terms relating to behavioural responses including Learning (GO:0007612), Social behaviour (GO:0035176) and Feeding behaviour (GO:0007631). Further in the damselfish we see enriched term Lens development in camera-type eye (GO:0002088) indicative of potential adaptations in the visual system to the distinct visual environments of the Caribbean in comparison to the TEP.

### GO Semantic Clusters in the Tropical Eastern Pacific

In the TEP we also find overlap of terms through the clustering analyses, though these had less clear links to environmental adaptation. Shared clusters included “response to external stimuli”, “signalling responses”, “development” and metabolic processes such as “Response to chemical”, “Phosphate-containing compound metabolic process”, “Signal transduction” and “Protein activation cascade” (Figure 1; Supplementary Table S2). The cluster “Response to chemical” included terms potentially relating to environmental stressors: Response to chemical (GO:0042221) and Cellular response to reactive oxygen species (GO:0034614) in the damselfish and Response to glucose (GO:0009749) and Response to bronchodilator (GO:0097366) in the groupers. These enrichment terms, particularly those related to reactive oxygen species (ROS) in the damselfish and glucose response in the groupers, may indicate modulation of metabolic and oxidative pathways in response to environmental stressors. “Phosphate-containing compound metabolic process” predominantly contained terms related to cellular and metabolic processes; of interest in the damselfish may be inositol phosphate metabolic process (GO:0043647), a signalling network involved in metabolic responses including nutrient sensing, maintenance of energy stores and control of energy homeostasis (Tu-Sekine and Kim 2022). Finally, terms within shared clusters “Signal transduction” and “Protein activation cascade” show enrichment in a multitude of signalling responses and pathways involved in responding to external signals, environmental cues, and mechanical stimuli (Figure 1; Supplementary Table S2).

### Gene Family Evolution

Our analyses of gene family evolution using CAFE5 (Mendes et al. 2021) on orthogroups identified through OrthoFinder (Emms and Kelly 2019) revealed differences in the patterns of change between the damselfish and groupers, particularly in how change has occurred between ocean basins. Whilst in the damselfish we found fairly similar levels of gene expansion in the TEP and Caribbean geminate (TEP: 54, CAR: 94), in the groupers we find gene expansion overwhelmingly occurring in the Caribbean geminate with little expansion occurring in the TEP (TEP: 34, CAR: 169; Figure 2). In terms of gene family contractions, the TEP and Caribbean geminate in the damselfish also showed similar, low levels of contraction (TEP: 34, CAR: 21). Meanwhile in the groupers we see a broad range of contractions in the TEP (466), while limited numbers are seen in the Caribbean (80). As with our analyses of selection, we found minimal parallelism in the functional enrichment of gene families found to be expanded or contracted between the groupers and damselfish. We therefore undertook the same clustering analysis approach to uncover broader similarities in functional change within the Caribbean and TEP.

**Figure 2.**
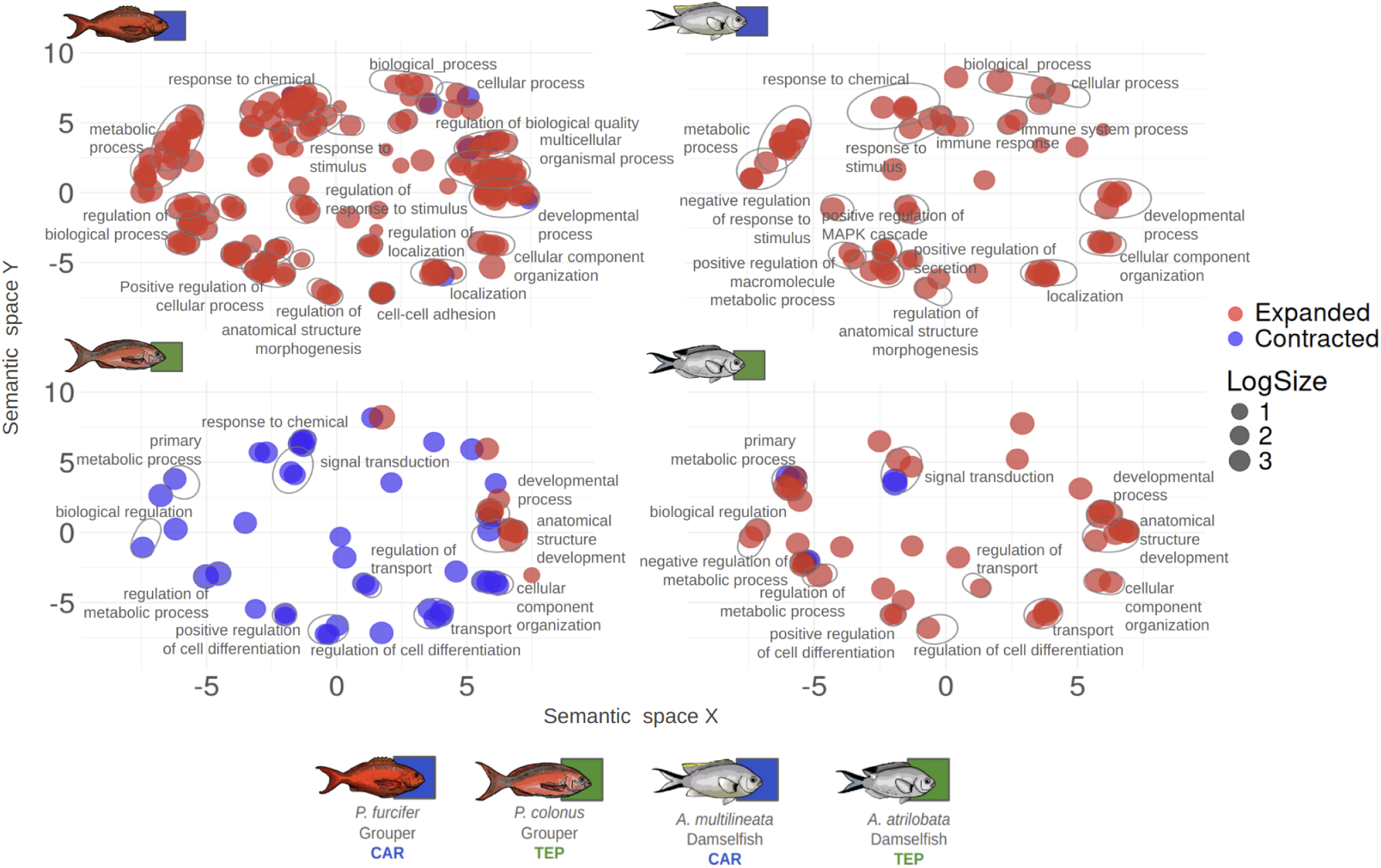
MDS plot of the pairwise semantic similarity of significantly enriched GO terms based on CAFE5 gene family analyses. More semantically similar GO terms will be closer together. Clustering analysis was performed on the two TEP species *A. atrilobata* (damselfish) and *C. colonus* (grouper) and on the two Caribbean species *A. multilineata* (damselfish) and *P. furcifer* (grouper) to identify clusters of similar GO terms between families in the same ocean environment. Clusters are identified with an ellipse and cluster label of the closest common ancestor GO term based on the GO hierarchy. LogSize refers to the frequency of the GO term in the underlying EBI Gene Ontology Annotation (GOA) database.

### GO Semantic Clusters in the Caribbean

In the Caribbean species of the damselfish and groupers, functional enrichment of gene family expansion revealed five terms functionally enriched in both families: DNA recombination (GO:0006310), Purine ribonucleotide metabolic process (GO:0009150), Organonitrogen compound biosynthetic process (GO:1901566), Organic hydroxy compound metabolic process (GO:1901615) and Positive regulation of MAPK cascade (GO:0043410) (Figure 2; Supplementary Table S3). Most of these overlapping terms are indicative of general fundamental metabolic and cellular processes; however, we again find the MAPK cascade as a shared process in this case expanding in both families. Since it is known to be involved in cellular responses to external environmental stressors such as UV radiation, salinity and chemical exposure, as well as regulating immune response, this pathway is likely relevant to adaptation to fluctuating environmental conditions (Cowan and Storey 2003, Liu et al. 2007).

Within the MAPK cascade cluster, we also see term Positive regulation of immune response (GO:0050778) in the damselfish and Positive regulation of phosphatidylinositol 3-kinase/protein kinase B signal transduction (GO:0051897) in the groupers. The PI3K-AKT signalling pathway has been indicated in stress response specifically being induced by exposure to low salinity (Cui et al. 2020). Also expanded, but not clustered, is the linked term positive regulation of MAP kinase activity (GO:0043406) in the grouper species.

We also identified a significant number of enriched clusters related to immune responses in both the groupers and the damselfish. Clusters such as “Response to stimulus”, “Regulation of response to stimulus”, “Immune response”, and “Immune system process” predominantly contained terms associated with immunity (Figure 2; Supplementary Table S3). For the groupers, relevant terms include Defense response to bacterium (GO:0042742), Humoral immune response (GO:0006959), Immune effector process (GO:0002252), Leukocyte migration (GO:0050900), and Leukocyte cell-cell adhesion (GO:0007159). In contrast, the damselfish displayed an even broader array of immune-related terms, including Inflammatory response (GO:0006954), Response to bacterium (GO:0009617), Leukocyte chemotaxis (GO:0030595), Regulation of immune response (GO:0050776), Regulation of defense response (GO:0031347), Adaptive immune response (GO:0002250), Innate immune response (GO:0045087), Leukocyte mediated immunity (GO:0002443), Myeloid leukocyte migration (GO:0097529), and Positive regulation of immune system process (GO:0002684). Additionally, several unclustered terms related to immune processes were present in the damselfish, including T cell proliferation (GO:0042098), Regulation of leukocyte activation (GO:0002694), Regulation of leukocyte proliferation (GO:0070663), and Interleukin-1 beta production (GO:0032611). Collectively, these terms underscore expansion of immune responses, tissue repair, and the organism’s capacity to defend against pathogens. Expansion of families enriched in such terms may be beneficial in environments characterised by high pathogen loads or frequent injuries, such as those particularly influenced by predators or pollution. Alternatively, since temperature is a major environmental modulator of immune systems in teleosts (Scharsack and Franke 2022), the influence of temperature on immune functions could also be relevant to the gene family expansions observed in these fish.

Whilst there is some overlap in the clustering of GO terms, in the Caribbean we also see multiple clusters of functional enrichment of expanded gene families in the grouper species which are not present in the damselfish including “Negative regulation of metabolic process”, “Negative regulation of cellular process”, “Cell-cell adhesion”, “Regulation of localization” and “Multicellular organismal process” indicative of the distinct strategies in how groupers have diverged and adapted to the same environmental shifts (Figure 2; Supplementary Table S3). Interestingly one cluster of functionally enriched terms in the gene family expansion found in the caribbean grouper includes the GO terms Feeding behavior (GO:0007631), Learning (GO:0007612), Behavior (GO:0007610) and Learning or memory (GO:0007611). This expansion tightly corresponds to a cluster of functionally enriched behaviour related terms in the selection analyses also in the Caribbean grouper.

Additionally, as in the selection analyses, in the groupers we found multiple clusters relevant to dealing with external stimuli including “Response to chemical”, “Response to stimulus”, “Regulation of response to stimulus” and “Negative regulation of response to stimulus”. These clusters are made up predominantly of expanded terms. Notably, those most indicative of environmental adaptation include Cellular response to starvation (GO:0009267), Cellular response to nutrient (GO:0031670), and Response to nutrient levels (GO:0031667). These terms highlight the fine-tuning of metabolic processes in response to fluctuations in nutrient availability. Additionally, Cellular response to ionizing radiation (GO:0071479) and Response to light stimulus (GO:0009416) may reflect requirement to adapt to the high light exposure in the oligotrophic Caribbean waters, as opposed to the more turbid waters of the TEP. Finally, another set of prominent terms include Cellular response to organic substance (GO:0071310), Response to organic substance (GO:0010033), Response to acid chemical (GO:0001101), Response to vitamin (GO:0033273), Xenobiotic metabolic process (GO:0006805), and Response to toxic substance (GO:0009636) and nearby, but not directly clustered term response to hypoxia (GO:0001666). These terms suggest that the groupers have expanded gene families relating to their capacity to manage and detoxify chemical compounds, possibly reflecting adaptations to anthropomorphic pollution or fluctuating chemical environments, including exposure to pollutants or naturally occurring toxins as well as hypoxic zones in the Caribbean.

### GO Semantic Clusters in the Tropical Eastern Pacific

In the TEP geminate species we see the biggest difference in the pattern of response between the damselfish and groupers. In the grouper geminate there is overwhelming contraction of gene families reflected in the functional enrichment, whereas in the damselfish we see a pattern of predominantly expansion (Figure 2). Further, we again see a limited number of functionally enriched terms shared between the grouper and damselfish geminates, in this case predominantly in developmental categories: epithelium development (GO:0060429), animal organ morphogenesis (GO:0009887) and system development (GO:0048731) (Figure 2; Supplementary Table S4). We find the opposite pattern in one shared term, regulation of neurogenesis (GO:0050767), which is expanded in the damselfish geminate but contracted in the grouper geminate.

Within the “Response to chemical” cluster, only present in the groupers, we see a multitude of contracted enriched terms indicative of external environmental cues: response to organonitrogen compound (GO:0010243), response to metal ion (GO:0010038), response to organic cyclic compound (GO:0014070), response to oxygen-containing compound (GO:1901700) and response to xenobiotic stimulus (GO:0009410). These terms indicate a contraction in gene families relating to responses to various chemical compounds present in the environments, ranging from naturally occurring organic molecules to potentially harmful pollutants like metal ions or xenobiotics. In a corresponding cluster of terms in the Caribbean we see similar terms expanded the damselfish (e.g. cellular response to oxygen-containing compound (GO:1901701)) and more prominently in the grouper geminate (e.g response to acid chemical (GO:0001101), xenobiotic metabolic process (GO:0006805) and response to toxic substance (GO:0009636)). The pattern of contraction in the TEP species and corresponding expansion in the Caribbean species of the groupers may indicate differences in the chemical composition or pollutants present in the two oceans potentially resulting in the requirement of specific mechanisms to metabolize, detoxify, or adapt to the presence of these compounds in the Caribbean.

Similarly potentially reflecting divergent evolutionary pressures tied to their environments, we also see term response to radiation (GO:0009314) contracted in the TEP grouper and concurrently term cellular response to ionizing radiation (GO:0071479) expanded in the Caribbean grouper (Figure 2; Supplementary Table S4). The oligotrophic waters of the Caribbean, which allow for greater light penetration, may have driven an expansion in cellular mechanisms to handle increased exposure to ionizing radiation, such as UV light. In contrast, the more turbid waters of the TEP, which receive less light, may have reduced the selective pressure for such radiation response mechanisms, leading to a contraction of these functions.

In the TEP damselfish we also see expansion in broad clusters relating to “Positive regulation of cell differentiation”, “Negative regulation of cell differentiation”, “Transport”, “Cellular component organization”, “Anatomical structure development” and “Developmental process”. One expansion enriched term, regulation of ERK1 and ERK2 cascade (GO:0070372), indicates another utilisation of the MAPK pathway in these fish. ERK1 and ERK2 signalling is directly involved in adaptive immunity in response and resistance to bacterial infection in teleosts (Wei et al. 2020, Du et al. 2024).

## Discussion

Taking advantage of the repeated experiments of divergence and adaptation promoted by the rise of the Isthmus of Panama, we produced high-quality whole genome assemblies of multiple reef fish. This provided an opportunity to investigate the predictability of adaptive response across the genome in ecologically similar, but phylogenetically distant species exposed to the same change in environmental conditions. Our comparative analyses of the damselfish and grouper geminates revealed that, whilst both families exhibited evidence of adaptive changes relating to the distinct environments in the TEP and Caribbean, indicated in both gene family expansions and signals of selection in coding genes, the specific responses and mechanisms were distinct. Some of the shared functional groups showing differentiation included immune response, signalling pathways relating to external stressors such as MAPKs and processes relating to nutrient availability. However, rarely did parallel evolution extend to the gene level or even specific GO terms. Predominantly we found functional parallel evolution only when we considered clusters of GO terms based on semantic similarity. In contrast to systems where more closely related species or populations showed evidence of parallel adaptive changes even to the amino acid level (e.g. Musilova et al. 2021), our findings indicate that phylogenetic constraints in this system may lead to quite different evolutionary outcomes.

Whilst there are striking examples of molecular parallelism in very distant species faced with the same selective pressures (Manceau et al. 2010, Feldman et al. 2012, Projecto-Garcia et al. 2013, Foote et al. 2015, Greenway et al. 2020, van Thiel et al. 2022), in these cases often the genetic basis of the adaptive trait may come into play, as well as the constraints acting on a particular system and the nature of the selection present. There are instances where the specific constraints of a molecular target may lead to a reduced number of solutions making molecular parallelism more likely, even in distant lineages, as is the case in toxin resistance (Feldman et al. 2012, van Thiel et al. 2022) and visual pigments (Musilova et al. 2021). In other cases a multitude of additional factors are likely to be influential including whether there are deleterious pleiotropic consequences (Chevin et al. 2010, Projecto-Garcia et al. 2013), mutational biases (Chan et al. 2010, Storz 2016), even the size of the gene (Moran et al. 2023). Here, when looking across the whole genome at changes relating to shifting environmental conditions which are likely relevant and representative of those encountered by many species, and potentially mirroring those which we will continue to see with global environmental change, the predominant difference in molecular response indicates the flexibility of these genetic systems and how species with similar ecologies can respond to very similar environmental-based selection in very different ways.

Not only do we see differences in the gene targets of selection between the geminates, but also in the type of response. In the positive selection analysis we found increased hits between the damselfish geminates when compared with the groupers, but perhaps most striking is the difference in the patterns of gene expansion and contraction between the families. Whilst the damselfish showed relatively equal patterns of predominant expansion in both the TEP and Caribbean species, the pattern in the groupers was very different. There was predominant gene family expansion in the Caribbean species and overwhelming contraction in the TEP species. Interestingly sometimes these represented corresponding clusters of areas which are seemingly being concurrently decreased in the TEP and expanded in the Caribbean, potentially representing selective responses specific to environmental challenges present only in the Caribbean. These involved chemical stress responses linked to various pollutants and radiation, potentially reflecting the higher levels of UV radiation present in the oligotrophic Caribbean, as well as the potentially greater influence of anthropogenic pollutants in these waters, though the latter may represent too recent a stressor to account for these patterns. Significant gene family expansion and contraction has commonly been linked with ecological niche shifts or significant environmental shifts in other species. For example, in one successful species of freshwater unionid bivalves, gene family expansion, particularly of cytochrome P450 genes, has been found to be driving adaptive evolution to strong anthropogenic environmental challenges which are resulting in the decline of many other freshwater Unionidae (Rogers et al. 2021). In some lineages there is also evidence of certain consistent ecological transitions leading to predictable expansion and contractions in certain gene families (Vertacnik et al. 2021), though we did not find evidence of this between families here.

The differences observed among the two groups in both the analyses of selection and family expansion or contraction is likely to also be influenced by the differences in the timing of their separation. Our divergence time estimates were consistent with an earlier separation of the damselfish geminates compared with the groupers (Craig and Hastings 2007, Lessios 2008, Rabosky et al. 2018, McCord et al. 2021), likely before the full emergence of the isthmus. Given the gradual process of the rise of the isthmus, the timing of a final cessation of gene flow has commonly been found to be different between species (Knowlton and Weigt 1998, Marko 2002, Lessios 2008, Marko and Moran 2009, Miura et al. 2010). This earlier separation may have provided damselfish with a longer window for independent adaptation, helping to explain the distinct patterns of genomic divergence recovered.

One of the key environmental shifts between the Caribbean and TEP post-isthmus closure is the decline in upwellings and consequently productivity in the Caribbean, both of which remain key features of the TEP today (Jackson and O’Dea 2023). Our expectation that this distinct difference in nutrient availability would be a key selective difference between the two oceans appears to have been met, however interestingly the response was most prominent in the groupers, and the two families have responded in different ways. In the Caribbean grouper expanded gene families included a cluster of functionally enriched terms relating to nutrient availability. Contrastingly in the Caribbean damselfish we found no evidence of gene expansion in related terms, but instead a similar term “Cellular response to nutrient levels” (GO:0031669) enriched in the positive selection analyses. In this case we may be seeing the impact of the interaction between an environmental pressure we expect to be exerting selective pressure on these species (nutrient availability) and that of past phylogenetic constraints due to the historic differences in feeding ecology between these two families (specialised planktivores versus historic demersal predators), as well as differences in divergence time, leading to differences in response.

Despite the predominance of different molecular targets in response to the environmental pressures of the TEP versus Caribbean, the few cases of parallel changes at the gene or pathway level tend to correspond to areas we hypothesised to be under selection given the environmental changes in the two basins. In the Caribbean geminates, we found the same zona pellucida related gene to be under selection in both families. Its function in egg envelope structure and sperm binding may reflect differences in reproductive strategy between the two oceans. The expectation of differences in the level of provisioning depending on productivity regime have been met in multiple trans-ithmanian geminate pairs of echinoderms (Lessios, 1990), arcid bivalves (Moran, 2004) and reef fish (Robertson and Collin, 2015), where larger egg sizes were consistently found in species residing in the lower productivity waters of the Caribbean when compared with their TEP counterparts. Interestingly, shifts in gene family size were additionally found in a zona pellucida protein gene family in another geminate pair of damselfish species investigated (Tracy et al. 2025), making it a consistent finding in all three geminate species so far investigated at the genome level. Taken together, these findings raise the possibility that zona pellucida related reproductive genes may represent a consistent molecular target of adaptation to the contrasting reproductive environments of the TEP and Caribbean.

Another consistent target in the Caribbean geminates was the MAPK signal transduction pathways. These pathways have been implicated in stress response in a wide range of species (Sheikh-Hamad and Gustin 2004, Kyriakis and Avruch 2012, Lin et al. 2021) and in response to a range of environmental stressors (Cowan and Storey 2003). Increasing evidence is building for their similar role in teleosts, with various MAPK signalling pathways involved in responses to salinity stressors (Kültz and Avila 2001, Tian et al. 2019, Duan et al. 2022), cold stress (Chu et al. 2021, Yu et al. 2022), immune response to bacterial infection (Qiao et al. 2021, Zhang et al. 2023), hypoxic conditions (Zhou et al. 2020, Yu et al. 2021) and environmental pollutants (Silvestre 2020). Specifically several MAPK pathways have been found to play a role in salinity adaptation as part of the osmosensory signalling pathways in the gills of Killifish, *Fundulus heteroclitus* (Kültz and Avila 2001) and identified in genome-wide association analysis in the nile tilapia (*Oreochromis niloticus*) in adaptation to oxygen stress, with MAPK signalling involved in the later growth stage under the hypoxic environment (Yu et al. 2021).

Specific MAPK gene *MAP3K10* is part of the ERK subfamily and was picked up in the Caribbean geminates of both the groupers and damselfish as under positive selection. Furthermore, in our gene enrichment analysis of selected genes we also found MAPK related terms in both families: “Regulation of p38MAPK cascade” (GO:1900744) in the damselfish and “Positive regulation of MAP kinase activity” (GO:0043406) in the groupers. Not only do we find shared signals of selection related to this signalling pathway, but also shared signals of gene family expansion with term “Positive regulation of MAPK cascade” (GO:0043410) one of the few functionally enriched terms found in both Caribbean species of the damselfish and groupers. Considering the higher temperature regimes and higher salinity levels in the Caribbean (D’Croz and Robertson 1997, Jackson and O’Dea 2023) and the evidence in other teleost species of the involvement of this pathway in controlling cellular responses to such environmental stressors we hypothesise this is a likely a candidate of a parallel adaptive response to the environmental conditions in the Caribbean between these geminates.

In the TEP evidence of parallel response came from different targets to that of the Caribbean. We found Solute carrier gene *SLC46A2* under selection in the TEP geminates of both the damselfish and groupers. Solute carriers (SLC) function in teleosts in exchange and transport of various molecules across membranes in maintenance of homeostasis across a range of physiological processes (Verri et al. 2012). In teleosts they demonstrate remarkable molecular diversity, adapting to species-specific physiological needs and environmental conditions (Romano et al. 2013). Expression patterns of certain SLC gene families have been found to vary in response to salinity and alkalinity stresses, indicating their importance in osmoregulation and acid-base homeostasis (Wang et al. 2020, Zhou et al. 2022). In particular, the gene *SLC46A2* has previously been found differentially expressed in response to salinity stress in transcriptomic studies of freshwater fish *Cottus rhenanus* (Escobar-Sierra and Lampert 2024a) and in the Coho Salmon (*Oncorhynchus kisutch*) post acclimation to salinity (Maryoung et al. 2015) indicating this gene as a potential adaptive target here in the two TEP geminates related to the maintenance of homeostasis in the dynamic conditions of the TEP.

As expected, we also see evidence of immune response being a target in both families with the same three immune related genes, *POU2AF1*, *TRIM16* and *SKAP1,* under positive selection in the TEP. Immune system function in fish, both innate and adaptive, can be particularly susceptible to various environmental stressors (Bly et al. 1997). A multitude of environmental factors including salinity, temperature, light intensity, oxygen levels, and pH can modulate and suppress immune responses in fish (Cuesta et al. 2007, Bowden 2008, Birrer et al. 2012, Choi et al. 2013, Maryoung et al. 2015, Lu et al. 2022, Franke et al. 2024). The immune system is also a target in the Caribbean species, with a shared expansion in different immune related terms in both Caribbean geminates though particularly strong in the damselfish where eleven enrichment terms showed explicit relation to immune function. Immune related genes have commonly been identified as a selective target in research investigating adaptive responses to the environment in natural populations of fish (Bernardi et al. 2016, Wilder et al. 2020, Escobar-Sierra and Lampert 2024a). Yet it is challenging to make direct predictions on the effects of adaptation in these groups, given the fast rate of evolution of immune genes across the tree of life (Kosiol et al. 2008, Hillier et al. 2004, Sackton et al. 2007, Lenz et al. 2013, Vinkler et al. 2023).

Whole-genome comparative studies in informative systems, such as the geminates considered in this study, provide a strong starting point from which to better understand the molecular targets of environmental selective pressures and the prevalence of adaptive molecular parallelism. However, it is important that parallel changes at the molecular level alone not be taken as evidence for local adaptation. Subsequent tests confirming the functional effects of putatively adaptive loci should be carried out in order to validate findings (Natarajan et al. 2016, Storz 2016). This will in turn enable us to better tease apart both the prevalence and contributing factors which may hinder or promote repeated adaptive solutions to similar environmental pressures in even distant lineages. Our findings of parallelism at the coarse functional level, but limited parallelism at the gene level may be relevant for better understanding evolutionary trajectories in marine species in response to changing environmental conditions. The six genomes sequenced, annotated and analyzed in this study represent a step forward in our understanding of the evolution of neotropical marine fishes, and are a valuable resource to study processes across the fish tree of life.

## Materials and Methods

### Sample Collection and Genomic Library Preparation

We collected adults of both our damselfish geminate species (*A. atrilobata* and *A. multilineata*) and grouper geminate species (*C. colonus* and *P. furcifer*) as well as closely related outgroups (*A. cyanea* for the damselfish and *C. fulva* for the groupers). Muscle tissue was immediately dissected and flash frozen on liquid nitrogen. Upon arrival in Panama City, we transferred tissue samples to -80°C for storage at the Smithsonian Tropical Research Institute (collection permit: SE/AO-4-19 issued by MiAmbiente). We extracted high molecular weight DNA from frozen muscle tissue using Qiagen Genomic-tip kits (Qiagen). We checked for the integrity of the DNA with a field inversion gel (Femto Pulse, Agilent). On average, DNA fragments were ∼50 kb in length. We prepared a PacBio HiFi library with the SMRTbell Express Template Prep Kit 2.0 (PacBio 100-938-900). We first sheared the DNA to 18 kbp using a Diagenode Megaruptor. We then selected fragments >15 kb with the BluePippin system (Sage Science), followed by AMPure bead cleanup. We sequenced the genomic library on a single 30-h movie 8M SMRT cells in CCS mode on the PacBio Sequel II system at the Brigham Young University DNA Sequencing Center.

### Genome Characterisation, Assembly and Annotation

We estimated genome size, heterozygosity, repeat and duplicate content of raw PacBio reads using GenomeScope (Vurture et al. 2017). We counted k-mers and generated a k-mer frequency distribution for 21-mers with Jellyfish v.2.2.6 (Marçais and Kingsford 2011) and the resulting histogram was later processed with GenomeScope. Raw genomic reads were assembled *de novo* using Hifiasm v. 0.16.1 (Cheng et al. 2021).

To identify and remove any contamination prior to annotation, we screened our genome assembly with taxon-annotated GC-coverage plots using BlobToolKit v.3.0 (Laetsch and Blaxter 2017, Challis et al. 2020). To prepare data for Blobtools, we mapped raw reads against the final genome assembly using minimap2 v2.28 (Li 2018). The resulting bam files were then sorted and merged with Samtools *sort* and *merge* commands (Danecek et al. 2021). A reference database for taxonomic assignment of contigs was created with MegaBLAST (Morgulis et al. 2008) using the following parameters: -task megablast and -e-value 1e-25. We used the Blobtools module map2cov to calculate coverage and generated a database with the Blobtools command *create.* Blobtools results were visualised and plotted with the BlobToolKit command *view*. Summary assembly statistics were estimated using Assembly_Stats v.0.14 (Trizna 2020). We ran Benchmarking Universal Single Copy Orthologs (BUSCO) v. 5.3 (Manni et al. 2021b, 2021a) to assess the completeness of the genomes. We scanned all the sequences for a ray-finned fish specific database of 3640 conserved genes (actinopterygii_odb10). Furthermore, we estimated assembly completeness and consensus quality value (QV) by counting kmers using meryl v1.3 (Rhie et al. 2020) with a *K*-value of 20 and inputting the meryl database, along with the final version of the assembly, to Merqury v1.3 (Rhie et al. 2020). Finally, we performed a synteny analysis with the D-GENIES web-based software (Cabanettes and Klopp 2018), using minimap2 for alignment. We further used D-GENIES to visualise synteny between geminate species and their outgroups.

We estimated the transposable element (TE) composition using RepeatModeler v.2.03 (Flynn et al. 2020) and RepeatMasker v.4.1.2 (Nishimura 2000) pipelines. We first constructed a *de novo* repeat library with RepeatModeler and we then annotated and masked the repeats in our final assemblies using RepeatMasker. We generated gene models using an automated pipeline (https://github.com/marchoeppner/genomeannotator, commit hash 1be8247). In this workflow, different types of evidence are processed and aligned against a repeat-masked assembly to inform the computation of a consensus gene build (https://github.com/marchoeppner/genomeannotator/blob/dev/CITATIONS.md). We used protein sequences from Uniprot, limited to Teleosts, and with support from either protein and/or transcriptome data -syntenically mapped gene models from two other fish species (*Maylandia zebra* and *Amphiprion percula*, EnsEMBL release 105) and species-specific transcriptome data where available. We estimated the completeness of the resulting gene builds using BUSCO against the actinopterygii_odb10 reference database. To functionally annotate each respective gene build, we analysed the final GFF3 file with EggnogMapper v2.1.9 (Cantalapiedra et al. 2021) against the core database v5.0.2 (Huerta-Cepas et al. 2019).

Additional to the reference genomes generated through this project, we utilised publicly available genomes of additional species for each family and annotated them using the same pipeline to ensure the annotation between species was as consistent as possible. For the groupers this consisted of *Epinephelus cyanopodus, Epinephelus fuscoguttatus, Epinephelus lanceolatus, Epinephelus moara, Centropristis striata, Cromileptes altivelis and Plectropomus leopardus.* For the damselfish we used *Acanthochromis polyacanthus, Amphiprion clarkii, Amphiprion ocellaris, Amphiprion percula and Dascyllus trimaculatus.* We additionally utilised genome assemblies of damselfish species *Abudefduf troschelii* and *Abudefduf saxatilis* whose annotation followed the same pipeline, details of which are outlined in Tracy et al. 2025. Accession numbers for the genomes and RNA-seq evidence data used are outlined in Supplementary Table S1.

### Positive Selection Analyses

To detect potential signatures of positive selection driving protein evolution in geminate species we used the branch-site model (Zhang et al. 2005) as configured in ETE-Toolkit v3.1.20 (Huerta-Cepas et al. 2016). As input we identified single copy orthologs between the geminate pair, outgroup and additional seven background species of the same family (ten species in total) for the damselfish and grouper comparisons using OrthoFinder v2.5.5 (Emms and Kelly 2019). Subsequent sets of orthologous gene sequences were aligned using the codon-aware model in PRANK v170427 (Löytynoja 2014). Using the ete-evol tool in ETE-Toolkit we estimated branch-site dN/ds ratios to identify positive selection at sites on particular foreground branches (Yang and Nielsen 2002, Zhang et al. 2005). For each family two runs were undertaken with either the TEP or Caribbean geminate marked as the foreground branch. Likelihood ratio tests (LRTs) were then applied to compare which of the models; a model of positive selection (bsA) or the null model of neutral evolution (bsA1), fit the data better. Genes were considered to show evidence of positive selection if the likelihood ratio test (LRT) was significant at *p* < 0.05, we also report adjusted *p*-values using the Benjamini-Hochberg procedure to control for multiple testing, with an FDR threshold of 10%.

### Molecular Divergence Analysis

To investigate molecular divergence across geminate species, we analysed 842 orthologous genes identified through OrthoFinder and confirmed to be shared between the families through shared annotation; genes showing evidence of positive selection in either lineage were excluded. All alignments were performed using MUSCLE in Geneious. We estimated Ka (non-synonymous substitution rate), Ks (synonymous substitution rate), the Ka/Ks ratio, and Dxy (nucleotide divergence per site). Ka and Ks values were calculated using KaKs_Calculator v2.0, while Dxy was computed using a custom script that determines the average number of nucleotide differences per site between sequences. Divergence times (in millions of years) were then estimated using the formula T = Ks / 2r, assuming a mutation rate of 1 × 10−8 substitutions per site per year and species-specific generation times: 2 years for the damselfish species and 5 years for the grouper species.

### Gene Family Evolution

To identify differences in gene family size dynamics between geminate species we employed CAFE5 v5.1.0 (Mendes et al. 2021) which estimates a birth-death parameter (λ) for a given tree and gene family counts. Our gene families were based on orthogroups identified by OrthoFinder (Orthogroups.GeneCount.tsv). Ultrametric phylogenetic trees, trimmed to include only the relevant species using R package Ape v5.8 (Paradis and Schliep 2019), were taken from McCord et al. (2021) for the damselfish and Rabosky et al. (2018) for the groupers. As recommended in the CAFE5 tutorial to avoid non-informative parameter estimates, families with particularly high variance in gene copy number were initially removed and subsequently analysed with the lambda estimate obtained from the remaining gene families. Prior to the final run of CAFE5 we calculated an appropriate error model to account for any errors present in the genome assemblies and trialled multiple values for the number of *gamma* rate categories until the log-likelihood score showed no improvement, finally selecting the best fit based on the log-likelihood score (Supplementary Table S6). Multiple runs were undertaken to ensure convergence. The *gamma* rate category for which we had the lowest log-likelihood did not converge following multiple runs, therefore the lowest log-likelihood for which we could get convergence was used, this was *gamma* = 2 in both families (Supplementary Tables S7-S8). We identified gene families significantly expanded or contracted in both TEP and Caribbean geminates based on *p*<0.05.

### GO Enrichment and Clustering

To identify gene functions over-represented in genes found to be under positive selection and gene families found to be undergoing significant expansion/contraction, we performed Gene Ontology (GO) enrichment analysis using Bioconductor package topGO v2.56.0 (Alexa and Rahnenfuhrer 2009) with the “Weight01” algorithm and Fisher’s exact test (*p*-value < 0.05). To improve the interpretability of GO terms found to be significantly enriched and facilitate broader scale comparison between the damselfish and groupers we used the REVIGO v1.8.1 core package (Supek et al. 2011) which clusters significant GO terms based on their semantic similarity using the SimRel similarity measure (Schlicker et al. 2006). Given the objective of this study to compare the enriched GO terms of different species, we modified the standard REVIGO workflow so that both families’ significant GO terms were analysed together, but without collapsing redundant terms between the two species.

The subsequent similarity matrix was run through the multidimensional scaling (MDS) procedure implemented in REVIGO and we used the MDS matrix as input into DBSCAN v1.1-12 (Hahsler et al. 2019) in order to formally identify clusters of similar GO terms. This enabled us to visualise key functional clusters of GO terms and identify overlapping clusters between the two families. In DBSCAN we set the minimum points parameter to 2 and set the “eps” parameter using a k-nearest neighbour distance plot and identifying the point at which there is a sudden increase in the k-nearest neighbour where we expect points beyond to be outliers (Hahsler et al. 2019). In order to more formally identify the “knee” point for the kNN-dist plots we employed the “Kneedle” algorithm, implemented in R package Kneedle v1.0.0 (Satopaa et al. 2011). Clusters were given an overall “descriptor” term label by tracing back in the GO ontology tree to identify the closest common ancestor term that covered at least 80% of the GO terms within a given cluster. We limited the clustering of GO terms to look at those only within the Biological Processes category as we felt this allows for more interpretable insights between genetic changes and how they might impact fitness or adaptation in comparison to molecular functions or cellular components.

