## Supplementary Information for "Comparative genomic analyses of trans-ithmanian reef fishes reveals different molecular targets of environmental adaptation in different families"

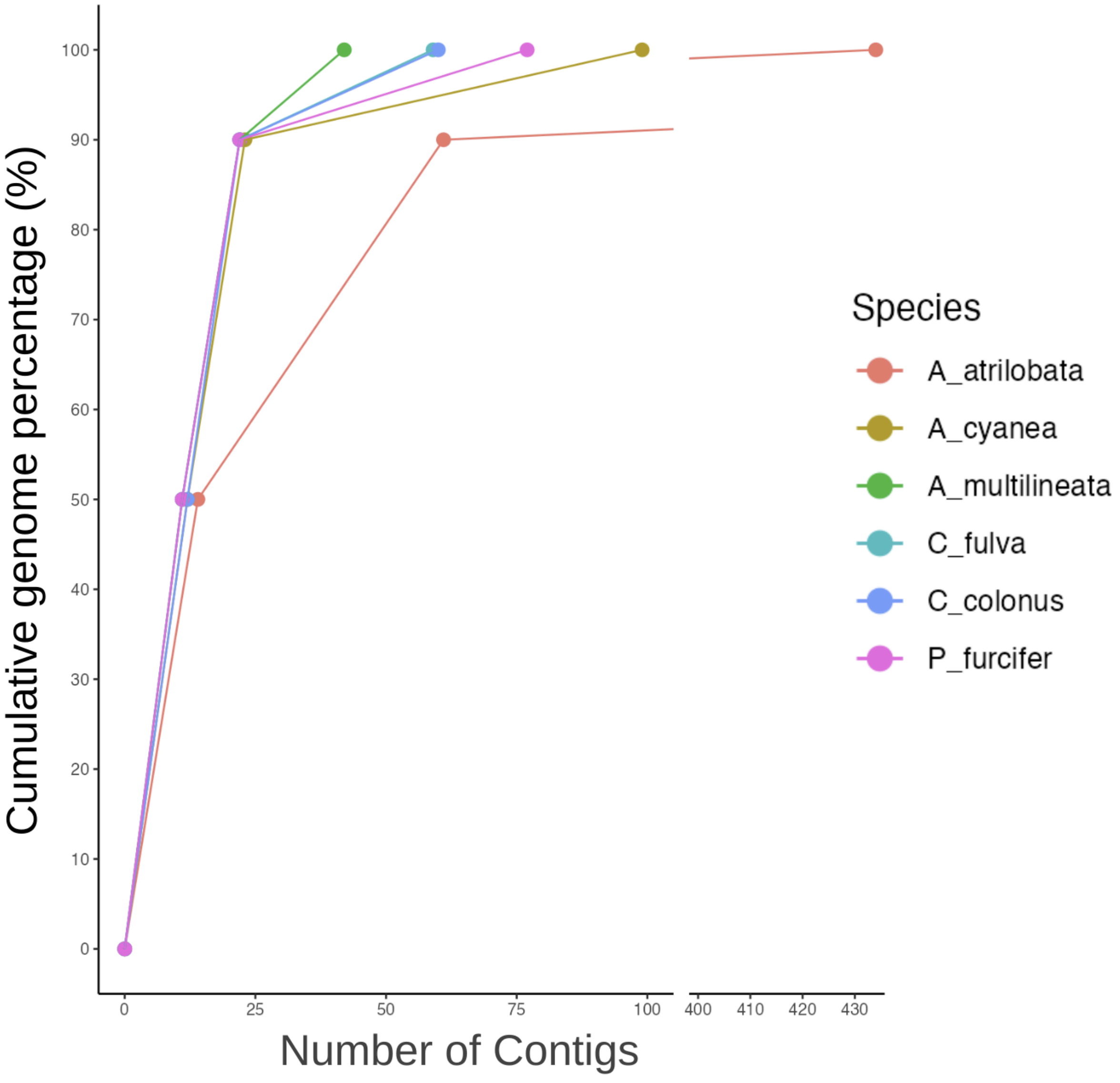


**Figure S1.** Contig number accumulation curves for the damselfish and grouper geminate species pairs and outgroups.

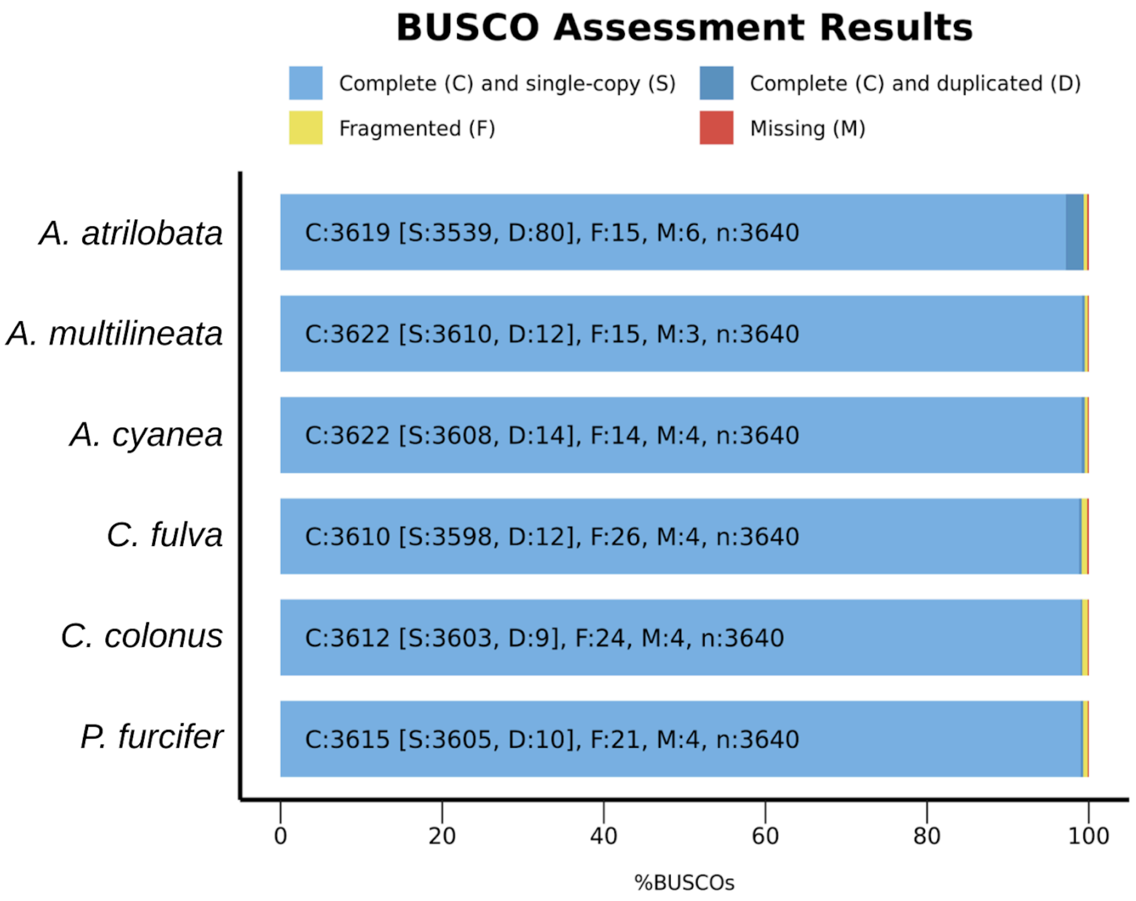


**Figure S2.** BUSCO assessment of the completeness of each species' genome assembly, benchmarked against the ray-finned fish specific database of 3640 conserved genes (actinopterygii_odb10).

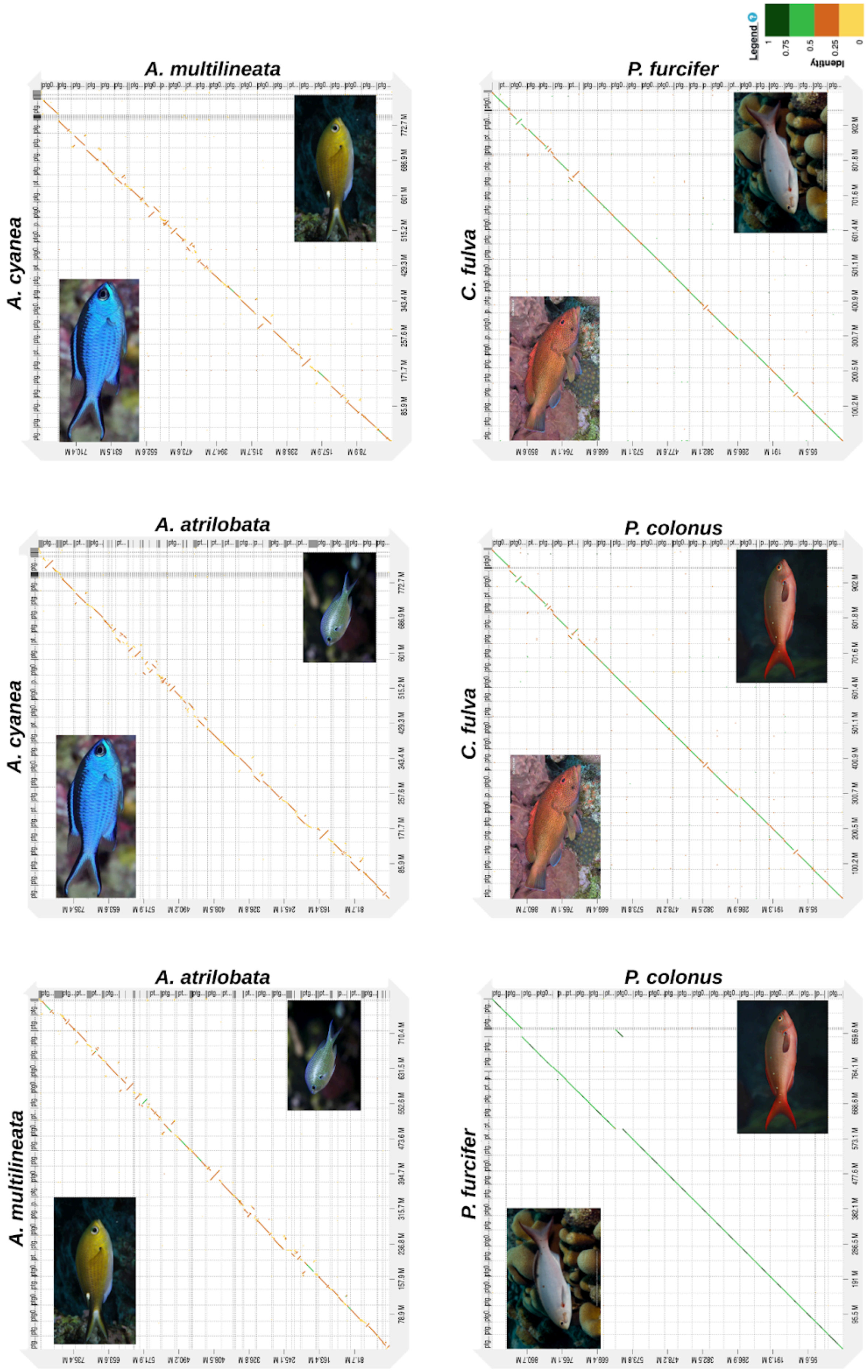


**Figure S3.** Conserved synteny analysis between geminate species and their outgroups; *A. atrilobata*,

*A. multilineata* and *A. cyanea* (outgroup) in the damselfish and *C. colonus*, *P. furcifer* and *C. fulva* (outgroup) in the groupers.

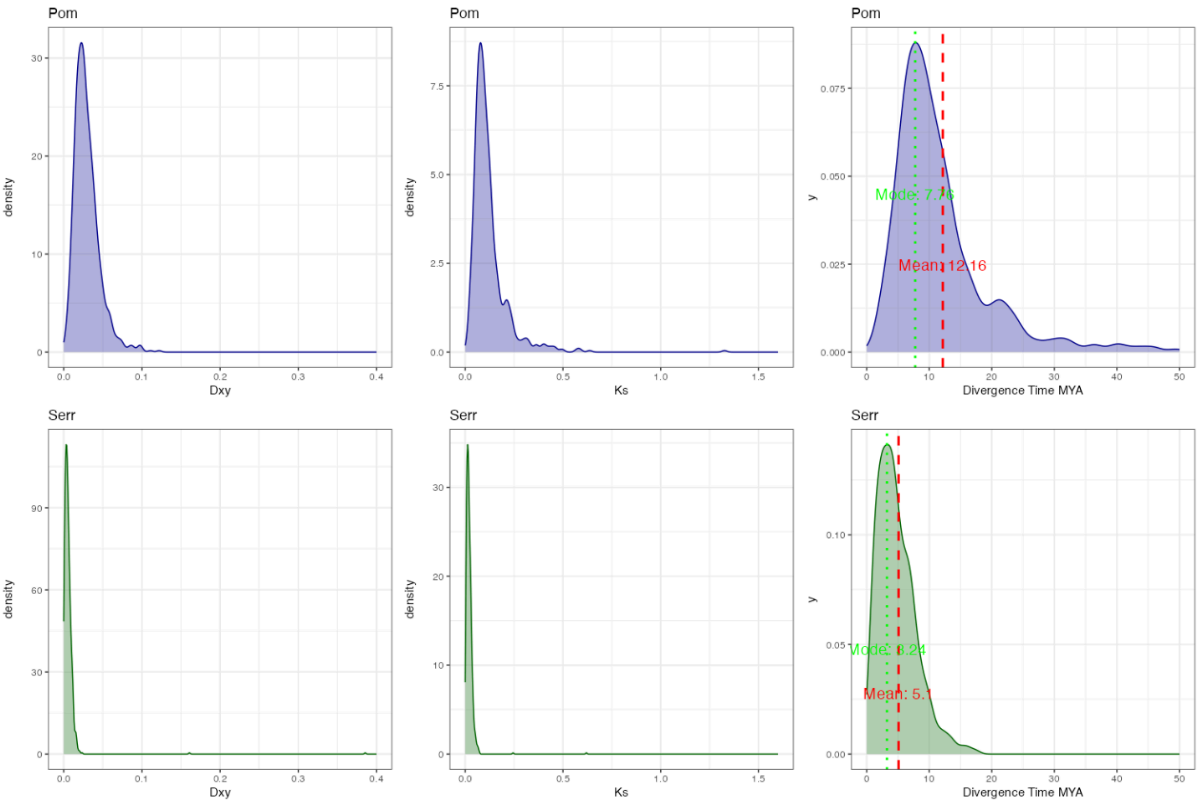


**Figure S4.** Genomic divergence metrics for the damselfish (*Pomacentridae*) and grouper (*Serranidae*) geminate species. Density plots show the distributions of Dxy, Ks and estimated divergence times (MYA) based on 842 orthologous genes. Each row corresponds to one species pair, with damselfish (Pom) in the top panels and grouper (Serr) in the bottom panels.

**Table S1.** Significantly enriched GO terms ($p$-value < 0.05) of genes identified through positive selection analyses in the Caribbean geminate of the damselfish *Azurina multilineata* and grouper *Paranthias furcifer*. Terms within each cluster identified in the MDS plot are listed. Cluster labels assigned to each cluster represent the closest common GO ancestor for that cluster. Shared GO terms between the damselfish and groupers are shown in bold.

| **Damselfish CAR geminate: *A.multilineata*** | | **Grouper CAR geminate: *P. furcifer*** | |
|---|---|---|---|
| **Custer: Cellular process** | | | |
| GO:0051674 | localization of cell | GO:0051649 | establishment of localization in cell |
| GO:0006898 | receptor-mediated endocytosis | GO:0060402 | calcium ion transport into cytosol |
| GO:0071702 | organic substance transport | GO:0043113 | receptor clustering |
| **Cluster: Developmental process** | | | |
| GO:0032989 | cellular component morphogenesis | GO:0001763 | morphogenesis of a branching structure |
| GO:0060560 | developmental growth involved in morphogenesis | GO:0000902 | cell morphogenesis |
| GO:0021510 | spinal cord development | GO:0007286 | spermatid development |
| GO:0001707 | mesoderm formation | - | - |
| GO:0030850 | prostate gland development | - | - |
| GO:0031100 | animal organ regeneration | - | - |
| GO:0003151 | outflow tract morphogenesis | - | - |
| GO:0048546 | digestive tract morphogenesis | - | - |
| GO:0001889 | liver development | - | - |
| GO:0003281 | ventricular septum development | - | - |
| GO:0048667 | cell morphogenesis involved in neuron differentiation | - | - |
| GO:0003208 | cardiac ventricle morphogenesis | - | - |
| GO:0031175 | neuron projection development | - | - |
| GO:0060351 | cartilage development involved in endochondral bone morphogenesis | - | - |
| GO:0035162 | embryonic hemopoiesis | - | - |
| GO:0002009 | morphogenesis of an epithelium | - | - |
| GO:0001947 | heart looping | - | - |
| GO:0002088 | lens development in camera-type eye | - | - |
| GO:0001541 | ovarian follicle development | - | - |
| GO:0055008 | cardiac muscle tissue morphogenesis | - | - |
| GO:0002040 | sprouting angiogenesis | - | - |
| GO:0048806 | genitalia development | - | - |

| Cluster: Positive regulation of biological process | | | |
|---|---|---|---|
| **GO:0045944** | **positive regulation of transcription by RNA polymerase II** | **GO:0045944** | **positive regulation of transcription by RNA polymerase II** |
| GO:0010628 | positive regulation of gene expression | GO:0051496 | positive regulation of stress fiber assembly |
| GO:0050679 | positive regulation of epithelial cell proliferation | - | - |
| GO:0051247 | positive regulation of protein metabolic process | - | - |
| GO:0010828 | positive regulation of glucose transmembrane transport | - | - |
| GO:1903039 | positive regulation of leukocyte cell-cell adhesion | - | - |
| GO:0033138 | positive regulation of peptidyl-serine phosphorylation | - | - |
| GO:0014911 | positive regulation of smooth muscle cell migration | - | - |
| GO:1900373 | positive regulation of purine nucleotide biosynthetic process | - | - |
| GO:0045923 | positive regulation of fatty acid metabolic process | - | - |
| **Cluster: Multicellular organismal process** | | | |
| GO:0008015 | blood circulation | GO:0007612 | learning |
| GO:0003014 | renal system process | GO:0007631 | feeding behavior |
| GO:0086003 | cardiac muscle cell contraction | GO:0035176 | social behavior |
| **Cluster: Positive regulation of striated muscle tissue development** | | | |
| GO:0045844 | positive regulation of striated muscle tissue development | - | - |
| GO:0045666 | positive regulation of neuron differentiation | - | - |
| GO:0055025 | positive regulation of cardiac muscle tissue development | - | - |
| **Cluster: Regulation of signal transduction** | | | |
| GO:0030510 | regulation of BMP signaling pathway | GO:0043949 | regulation of cAMP-mediated signaling |
| GO:0050680 | negative regulation of epithelial cell proliferation | GO:0045744 | negative regulation of G protein-coupled receptor signaling pathway |
| GO:1900744 | regulation of p38MAPK cascade | - | - |
| **Cluster: Cellular component organization or biogenesis** | | | |
| GO:0016043 | cellular component organization | GO:0001941 | postsynaptic membrane organization |
| GO:0060271 | cilium assembly | - | - |

| | | | |
|---|---|---|---|
| **Cluster: Response to stimulus** | | | |
| GO:0097066 | response to thyroid hormone | GO:0071277 | cellular response to calcium ion |
| GO:0060326 | cell chemotaxis | GO:0006954 | inflammatory response |
| GO:0071549 | cellular response to dexamethasone stimulus | GO:0034605 | cellular response to heat |
| GO:0044344 | cellular response to fibroblast growth factor stimulus | GO:0071364 | cellular response to epidermal growth factor stimulus |
| GO:0071363 | cellular response to growth factor stimulus | - | - |
| GO:0006952 | defense response | - | - |
| GO:0071466 | cellular response to xenobiotic stimulus | - | - |
| GO:0046677 | response to antibiotic | - | - |
| GO:0010035 | response to inorganic substance | - | - |
| **Cluster: Cellular process** | | | |
| GO:0032879 | regulation of localization | GO:0008037 | cell recognition |
| GO:0007018 | microtubule-based movement | - | - |
| GO:0022406 | membrane docking | - | - |
| GO:0003341 | cilium movement | - | - |
| **Cluster: Cellular response to stimulus** | | | |
| GO:0031669 | cellular response to nutrient levels | GO:0071214 | cellular response to abiotic stimulus |
| - | - | GO:0007200 | phospholipase C-activating G protein-coupled receptor signaling pathway |
| **Cluster: Regulation of multicellular organismal process** | | | |
| GO:0060964 | regulation of miRNA-mediated gene silencing | GO:0034103 | regulation of tissue remodeling |
| - | - | GO:0050795 | regulation of behavior |
| **Cluster: Organonitrogen compound metabolic process** | | | |
| GO:0046068 | cGMP metabolic process | GO:0009069 | serine family amino acid metabolic process |
| - | - | GO:0006508 | proteolysis |
| - | - | GO:0043967 | histone H4 acetylation |
| **Unclustered** | | | |
| GO:0010717 | regulation of epithelial to mesenchymal transition | GO:0070085 | glycosylation |
| GO:2000027 | regulation of animal organ morphogenesis | GO:0034405 | response to fluid shear stress |
| GO:0007267 | cell-cell signaling | GO:0051209 | release of sequestered calcium ion into cytosol |
| GO:0006694 | steroid biosynthetic process | GO:0051726 | regulation of cell cycle |
| GO:0035150 | regulation of tube size | GO:0043406 | positive regulation of MAP kinase |

| | | | |
|---|---|---|---|
| | | | activity |
| GO:0010975 | regulation of neuron projection development | GO:0043392 | negative regulation of DNA binding |
| GO:0042698 | ovulation cycle | GO:0048168 | regulation of neuronal synaptic plasticity |
| GO:0051145 | smooth muscle cell differentiation | - | - |
| GO:0034446 | substrate adhesion-dependent cell spreading | - | - |
| GO:0090087 | regulation of peptide transport | - | - |
| GO:0040011 | locomotion | - | - |
| GO:0017158 | regulation of calcium ion-dependent exocytosis | - | - |
| GO:0019933 | cAMP-mediated signaling | - | - |
| GO:0030224 | monocyte differentiation | - | - |
| GO:0043368 | positive T cell selection | - | - |
| GO:0086001 | cardiac muscle cell action potential | - | - |
| GO:0072210 | metanephric nephron development | - | - |
| GO:0051099 | positive regulation of binding | - | - |

**Table S2.** Significantly enriched GO terms ($p$-value < 0.05) of genes identified through positive selection analyses in the Tropical Eastern Pacific geminate of the damselfish *Azurina atrilobata* and grouper *Cephalopholis colonus*. Terms within each cluster identified in the MDS plot are listed. Cluster labels assigned to each cluster represent the closest common GO ancestor for that cluster. Shared GO terms between the damselfish and groupers are shown in bold.

| **Damselfish TEP geminate: *A. atrilobata*** | | **Grouper TEP geminate: *C. colonus*** | |
|---|---|---|---|
| **Custer: Response to chemical** | | | |
| GO:0042221 | response to chemical | GO:0034614 | cellular response to reactive oxygen species |
| GO:0009749 | response to glucose | GO:0097366 | response to bronchodilator |
| **Custer: Cell adhesion** | | | |
| GO:0033627 | cell adhesion mediated by integrin | GO:0007156 | homophilic cell adhesion via plasma membrane adhesion molecules |
| GO:0007160 | cell-matrix adhesion | - | - |
| GO:0007159 | leukocyte cell-cell adhesion | - | - |
| **Custer: Negative regulation of biological process** | | | |
| GO:0050686 | negative regulation of mRNA | GO:0045744 | negative regulation of G protein-coupled |

| | | | |
|---|---|---|---|
| | processing | | receptor signaling pathway |
| GO:0030512 | negative regulation of transforming growth factor beta receptor signaling pathway | GO:0010977 | negative regulation of neuron projection development |
| GO:1903845 | negative regulation of cellular response to transforming growth factor beta stimulus | GO:0035023 | regulation of Rho protein signal transduction |
| GO:0043409 | negative regulation of MAPK cascade | GO:0046580 | negative regulation of Ras protein signal transduction |
| GO:0007162 | negative regulation of cell adhesion | GO:0030336 | negative regulation of cell migration |
| **Custer: Transport** | | | |
| **GO:0051674** | **localization of cell** | **GO:0051674** | **localization of cell** |
| GO:0035725 | sodium ion transmembrane transport | GO:0034502 | protein localization to chromosome |
| GO:0046717 | acid secretion | - | - |
| GO:0006898 | receptor-mediated endocytosis | - | - |
| GO:0050658 | RNA transport | - | - |
| GO:0051938 | L-glutamate import | - | - |
| **Custer: Cellular component organization** | | | |
| GO:0007131 | reciprocal meiotic recombination | GO:0006338 | chromatin remodeling |
| GO:0007044 | cell-substrate junction assembly | GO:0006325 | chromatin organization |
| GO:0070206 | protein trimerization | GO:0034332 | adherens junction organization |
| GO:0046847 | filopodium assembly | GO:0051276 | chromosome organization |
| GO:0051262 | protein tetramerization | - | - |
| GO:0140013 | meiotic nuclear division | - | - |
| GO:0051260 | protein homooligomerization | - | - |
| **Custer: Multicellular organismal process** | | | |
| GO:0001503 | ossification | GO:0003014 | renal system process |
| GO:0050905 | neuromuscular process | - | - |
| GO:0003018 | vascular process in circulatory system | - | - |
| **Custer: Positive regulation of biological process** | | | |
| **GO:0045070** | **positive regulation of viral genome replication** | **GO:0045070** | **positive regulation of viral genome replication** |
| GO:0010638 | positive regulation of organelle organization | GO:0050769 | positive regulation of neurogenesis |
| GO:0050731 | positive regulation of peptidyl-tyrosine phosphorylation | GO:0010800 | positive regulation of peptidyl-threonine phosphorylation |
| GO:0045944 | positive regulation of transcription by RNA polymerase II | GO:1903902 | positive regulation of viral life cycle |
| GO:0030858 | positive regulation of epithelial cell differentiation | GO:0051590 | positive regulation of neurotransmitter transport |

| | | | |
|---|---|---|---|
| GO:0032024 | positive regulation of insulin secretion | GO:0051155 | positive regulation of striated muscle cell differentiation |
| GO:0010628 | positive regulation of gene expression | GO:0045666 | positive regulation of neuron differentiation |
| GO:0048584 | positive regulation of response to stimulus | GO:0045840 | positive regulation of mitotic nuclear division |
| GO:0048522 | positive regulation of cellular process | - | - |
| GO:0010811 | positive regulation of cell-substrate adhesion | - | - |
| GO:0050850 | positive regulation of calcium-mediated signaling | - | - |
| GO:0001954 | positive regulation of cell-matrix adhesion | - | - |
| GO:0043525 | positive regulation of neuron apoptotic process | - | - |
| GO:0046638 | positive regulation of alpha-beta T cell differentiation | - | - |
| GO:0045684 | positive regulation of epidermis development | - | - |
| **Custer: Signal transduction** | | | |
| **GO:0007229** | **integrin-mediated signaling pathway** | **GO:0007229** | **integrin-mediated signaling pathway** |
| GO:0007224 | smoothened signaling pathway | GO:0035556 | intracellular signal transduction |
| - | - | GO:0007193 | adenylate cyclase-inhibiting G protein-coupled receptor signaling pathway |
| - | - | GO:0007200 | phospholipase C-activating G protein-coupled receptor signaling pathway |
| - | - | GO:0007187 | G protein-coupled receptor signaling pathway, coupled to cyclic nucleotide second messenger |
| **Custer: Phosphate-containing compound metabolic process** | | | |
| GO:0006195 | purine nucleotide catabolic process | GO:0042417 | dopamine metabolic process |
| GO:0008033 | tRNA processing | GO:0009179 | purine ribonucleoside diphosphate metabolic process |
| GO:0043647 | inositol phosphate metabolic process | GO:0044270 | cellular nitrogen compound catabolic process |
| - | - | GO:0046700 | heterocycle catabolic process |
| **Custer: Anatomical structure development** | | | |
| GO:0071695 | anatomical structure maturation | GO:0009880 | embryonic pattern specification |
| GO:0061383 | trabecula morphogenesis | GO:0001763 | morphogenesis of a branching structure |
| GO:0048705 | skeletal system morphogenesis | GO:0035987 | endodermal cell differentiation |
| GO:0000768 | syncytium formation by plasma membrane fusion | GO:0048813 | dendrite morphogenesis |

| | | | |
|---|---|---|---|
| GO:0014037 | Schwann cell differentiation | GO:0014003 | oligodendrocyte development |
| GO:0021766 | hippocampus development | GO:0001764 | neuron migration |
| GO:0048646 | anatomical structure formation involved in morphogenesis | GO:0021532 | neural tube patterning |
| - | - | GO:0021510 | spinal cord development |
| - | - | GO:0007566 | embryo implantation |
| **Custer: Protein activation cascade** | | | |
| GO:0072376 | protein activation cascade | GO:0043967 | histone H4 acetylation |
| GO:0035601 | protein deacylation | - | - |
| **Custer: Negative regulation of developmental process** | | | |
| GO:0045668 | negative regulation of osteoblast differentiation | GO:0010771 | negative regulation of cell morphogenesis involved in differentiation |
| - | - | GO:0051961 | negative regulation of nervous system development |
| **Unclustered** | | | |
| **GO:0007059** | **chromosome segregation** | **GO:0007059** | **chromosome segregation** |
| GO:0051445 | regulation of meiotic cell cycle | GO:0007416 | synapse assembly |
| GO:0007140 | male meiotic nuclear division | GO:0040011 | locomotion |
| GO:0016064 | immunoglobulin mediated immune response | GO:0097164 | ammonium ion metabolic process |
| GO:0051304 | chromosome separation | GO:0031279 | regulation of cyclase activity |
| GO:0044774 | mitotic DNA integrity checkpoint signaling | GO:0051339 | regulation of lyase activity |
| GO:0030224 | monocyte differentiation | GO:2000027 | regulation of animal organ morphogenesis |
| GO:0072676 | lymphocyte migration | GO:0048519 | negative regulation of biological process |
| GO:0036297 | interstrand cross-link repair | GO:0060218 | hematopoietic stem cell differentiation |
| GO:0042060 | wound healing | GO:0014910 | regulation of smooth muscle cell migration |
| GO:0071560 | cellular response to transforming growth factor beta stimulus | GO:0007204 | positive regulation of cytosolic calcium ion concentration |
| GO:0070613 | regulation of protein processing | - | - |
| GO:0051783 | regulation of nuclear division | - | - |
| GO:0009566 | fertilization | - | - |

**Table S3.** Significantly enriched GO terms (*p*-value < 0.05) identified through gene family expansion/contraction analyses in the Caribbean geminate of the damselfish *Azurina multilineata* and groupers *Paranthias furcifer*. Enriched expanded and contracted terms are listed by cluster as identified through semantic similarity. Cluster labels assigned to each cluster represent the closest common GO ancestor for that cluster. Expanded enrichment terms are shown with a (+) and contracted a (-). Shared GO terms between the damselfish and groupers are shown in bold.

| **Damselfish CAR geminate: *A. multilineata*** | | | **Grouper CAR geminate: *P. furcifer*** | | |
|---|---|---|---|---|---|
| **Custer: Response to chemical** | | | | | |
| GO:1901701 | cellular response to oxygen-containing compound | + | GO:0001101 | response to acid chemical | + |
| GO:0071495 | cellular response to endogenous stimulus | + | GO:0043434 | response to peptide hormone | + |
| GO:0071407 | cellular response to organic cyclic compound | + | GO:0043200 | response to amino acid | + |
| - | - | | GO:0071479 | cellular response to ionizing radiation | + |
| - | - | | GO:0071375 | cellular response to peptide hormone stimulus | + |
| - | - | | GO:0019221 | cytokine-mediated signaling pathway | + |
| - | - | | GO:0071345 | cellular response to cytokine stimulus | + |
| - | - | | GO:0031670 | cellular response to nutrient | + |
| - | - | | GO:0006805 | xenobiotic metabolic process | + |
| - | - | | GO:0010033 | response to organic substance | + |
| - | - | | GO:0009636 | response to toxic substance | + |
| - | - | | GO:0070848 | response to growth factor | + |
| - | - | | GO:0046677 | response to antibiotic | + |
| - | - | | GO:0033273 | response to vitamin | + |
| - | - | | GO:0033209 | tumor necrosis factor-mediated signaling pathway | + |
| - | - | | GO:0045471 | response to ethanol | + |
| - | - | | GO:0071310 | cellular response to organic substance | + |
| - | - | | GO:0032570 | response to progesterone | + |

| | | | | | |
|---|---|---|---|---|---|
| - | - | | GO:0009416 | response to light stimulus | + |
| - | - | | GO:0032570 | response to progesterone | - |
| **Custer: Response to stimulus** | | | | | |
| GO:0006954 | inflammatory response | + | GO:0042060 | wound healing | + |
| GO:0009617 | response to bacterium | + | GO:0009267 | cellular response to starvation | + |
| GO:0030595 | leukocyte chemotaxis | + | GO:0031667 | response to nutrient levels | + |
| - | - | | GO:0042742 | defense response to bacterium | + |
| **Custer: Regulation of response to stimulus** | | | | | |
| GO:0050776 | regulation of immune response | + | GO:0010469 | regulation of signaling receptor activity | + |
| GO:0031347 | regulation of defense response | + | GO:0046578 | regulation of Ras protein signal transduction | + |
| - | - | | GO:0035023 | regulation of Rho protein signal transduction | + |
| **Custer: Negative regulation of response to stimulus** | | | | | |
| GO:0048585 | negative regulation of response to stimulus | + | GO:1902532 | negative regulation of intracellular signal transduction | + |
| - | - | | GO:2001240 | negative regulation of extrinsic apoptotic signaling pathway in absence of ligand | + |
| - | - | | GO:1903170 | negative regulation of calcium ion transmembrane transport | + |
| **Cluster: Immune response** | | | | | |
| GO:0002250 | adaptive immune response | + | GO:0006959 | humoral immune response | + |
| GO:0045087 | innate immune response | + | - | - | |
| **Custer: Immune system process** | | | | | |
| GO:0002443 | leukocyte mediated immunity | + | GO:0002252 | immune effector process | + |
| GO:0097529 | myeloid leukocyte migration | + | GO:0050900 | leukocyte migration | + |
| **Custer: Metabolic process** | | | | | |
| **GO:0006310** | **DNA recombination** | + | **GO:0006310** | **DNA recombination** | + |
| **GO:0009150** | **purine ribonucleotide metabolic process** | + | **GO:0009150** | **purine ribonucleotide metabolic process** | + |
| **GO:1901566** | **organonitrogen compound biosynthetic process** | + | **GO:1901566** | **organonitrogen compound biosynthetic process** | + |
| GO:0036211 | protein modification process | + | GO:0044237 | cellular metabolic process | + |
| GO:0034654 | nucleobase-containing compound biosynthetic process | + | GO:1901565 | organonitrogen compound catabolic process | + |

| | | | | | |
|---|---|---|---|---|---|
| GO:0090407 | organophosphate biosynthetic process | + | GO:0046434 | organophosphate catabolic process | + |
| - | - | | GO:0006261 | DNA-templated DNA replication | + |
| - | - | | GO:0046496 | nicotinamide nucleotide metabolic process | + |
| - | - | | GO:0046474 | glycerophospholipid biosynthetic process | + |
| - | - | | GO:0006790 | sulfur compound metabolic process | + |
| - | - | | GO:0006468 | protein phosphorylation | + |
| - | - | | GO:0030258 | lipid modification | + |
| **Custer: Metabolic process** | | | | | |
| **GO:1901615** | **organic hydroxy compound metabolic process** | + | **GO:1901615** | **organic hydroxy compound metabolic process** | + |
| GO:1901137 | carbohydrate derivative biosynthetic process | + | GO:0009056 | catabolic process | + |
| - | - | | GO:1901136 | carbohydrate derivative catabolic process | + |
| - | - | | GO:0044282 | small molecule catabolic process | + |
| - | - | | GO:1901657 | glycosyl compound metabolic process | + |
| **Custer: Negative regulation of metabolic process** | | | | | |
| - | - | | GO:0031330 | negative regulation of cellular catabolic process | + |
| - | - | | GO:0010629 | negative regulation of gene expression | + |
| - | - | | GO:0031397 | negative regulation of protein ubiquitination | + |
| - | - | | GO:0009895 | negative regulation of catabolic process | + |
| - | - | | GO:0001818 | negative regulation of cytokine production | + |
| - | - | | GO:0051053 | negative regulation of DNA metabolic process | + |
| - | - | | GO:0000122 | negative regulation of transcription by RNA polymerase II | + |
| - | - | | GO:0051248 | negative regulation of protein metabolic process | + |
| **Custer: Positive regulation of macromolecule metabolic process** | | | | | |
| GO:0001819 | positive regulation of cytokine production | + | GO:0032147 | activation of protein kinase activity | + |
| GO:0045860 | positive regulation of protein kinase activity | + | GO:0045944 | positive regulation of transcription by RNA polymerase II | + |
| - | - | | GO:0010628 | positive regulation of gene expression | + |

| | | | | | |
|---|---|---|---|---|---|
| - | - | | GO:0050731 | positive regulation of peptidyl-tyrosine phosphorylation | + |
| - | - | | GO:0009891 | positive regulation of biosynthetic process | + |
| - | - | | GO:0071902 | positive regulation of protein serine/threonine kinase activity | + |
| - | - | | GO:0045732 | positive regulation of protein catabolic process | + |
| **Custer: Regulation of biological process** | | | | | |
| - | - | | GO:0042176 | regulation of protein catabolic process | + |
| - | - | | GO:0019216 | regulation of lipid metabolic process | + |
| - | - | | GO:2000377 | regulation of reactive oxygen species metabolic process | + |
| - | - | | GO:0006275 | regulation of DNA replication | + |
| - | - | | GO:0010506 | regulation of autophagy | + |
| **Custer: Positive regulation of cellular process** | | | | | |
| GO:0045785 | positive regulation of cell adhesion | + | GO:0030335 | positive regulation of cell migration | + |
| GO:0051094 | positive regulation of developmental process | + | GO:0010976 | positive regulation of neuron projection development | + |
| GO:0002684 | positive regulation of immune system process | + | GO:0045666 | positive regulation of neuron differentiation | + |
| GO:0050867 | positive regulation of cell activation | + | GO:0050769 | positive regulation of neurogenesis | + |
| - | - | | GO:0030307 | positive regulation of cell growth | + |
| - | - | | GO:0051781 | positive regulation of cell division | + |
| - | - | | GO:0050679 | positive regulation of epithelial cell proliferation | + |
| - | - | | GO:0090068 | positive regulation of cell cycle process | + |
| - | - | | GO:0045931 | positive regulation of mitotic cell cycle | + |
| - | - | | GO:0045787 | positive regulation of cell cycle | + |
| - | - | | GO:0010811 | positive regulation of cell-substrate adhesion | + |
| - | - | | GO:0022409 | positive regulation of cell-cell adhesion | + |
| **Custer: Negative regulation of cellular process** | | | | | |
| - | - | | GO:0043524 | negative regulation of neuron apoptotic process | + |
| - | - | | GO:0008285 | negative regulation of cell population proliferation | + |

| | | | | | |
|---|---|---|---|---|---|
| - | - | | GO:2000134 | negative regulation of G1/S transition of mitotic cell cycle | + |
| - | - | | GO:0031345 | negative regulation of cell projection organization | + |
| - | - | | GO:0050680 | negative regulation of epithelial cell proliferation | + |
| **Custer: Positive regulation of secretion** | | | | | |
| GO:0050714 | positive regulation of protein secretion | + | GO:1900182 | positive regulation of protein localization to nucleus | + |
| GO:0002793 | positive regulation of peptide secretion | + | - | - | |
| **Custer: Positive regulation of MAPK cascade** | | | | | |
| **GO:0043410** | **positive regulation of MAPK cascade** | + | **GO:0043410** | **positive regulation of MAPK cascade** | + |
| GO:0050778 | positive regulation of immune response | + | GO:0051897 | positive regulation of phosphatidylinositol 3-kinase/protein kinase B signal transduction | + |
| **Custer: Cell-cell adhesion** | | | | | |
| - | - | | GO:0098742 | cell-cell adhesion via plasma-membrane adhesion molecules | - |
| - | - | | GO:0007156 | homophilic cell adhesion via plasma membrane adhesion molecules | + |
| - | - | | GO:0007160 | cell-matrix adhesion | + |
| - | - | | GO:0034113 | heterotypic cell-cell adhesion | + |
| - | - | | GO:0007159 | leukocyte cell-cell adhesion | + |
| - | - | | GO:0098609 | cell-cell adhesion | + |
| **Custer: Regulation of anatomical structure morphogenesis** | | | | | |
| GO:2000026 | regulation of multicellular organismal development | + | GO:2000027 | regulation of animal organ morphogenesis | + |
| - | - | | GO:0045765 | regulation of angiogenesis | + |
| - | - | | GO:0022604 | regulation of cell morphogenesis | + |
| **Custer: Localization** | | | | | |
| GO:0071702 | organic substance transport | + | GO:0051674 | localization of cell | + |
| GO:0046907 | intracellular transport | + | GO:0002576 | platelet degranulation | + |
| GO:0051640 | organelle localization | + | GO:0008104 | protein localization | + |
| GO:0006897 | endocytosis | + | GO:0006810 | transport | + |
| GO:0071705 | nitrogen compound transport | + | GO:0051649 | establishment of localization in cell | + |

| | | | | | |
|---|---|---|---|---|---|
| - | - | | GO:0016192 | vesicle-mediated transport | + |
| - | - | | GO:1902476 | chloride transmembrane transport | - |
| **Custer: Regulation of localization** | | | | | |
| - | - | | GO:0090087 | regulation of peptide transport | + |
| - | - | | GO:0017158 | regulation of calcium ion-dependent exocytosis | + |
| - | - | | GO:1905475 | regulation of protein localization to membrane | + |
| **Custer: Cellular component organization** | | | | | |
| GO:0070925 | organelle assembly | + | GO:0030198 | extracellular matrix organization | + |
| GO:0034330 | cell junction organization | + | GO:0007043 | cell-cell junction assembly | + |
| GO:0022607 | cellular component assembly | + | GO:0034332 | adherens junction organization | + |
| - | - | | GO:0051260 | protein homooligomerization | + |
| **Custer: Celluar process** | | | | | |
| GO:0007018 | microtubule-based movement | + | GO:0008283 | cell population proliferation | + |
| - | - | | GO:0007623 | circadian rhythm | - |
| **Custer: Regulation of biological quality** | | | | | |
| GO:0007204 | positive regulation of cytosolic calcium ion concentration | + | GO:0042445 | hormone metabolic process | + |
| - | - | | GO:0050878 | regulation of body fluid levels | - |
| **Custer: Developmental process** | | | | | |
| GO:0032989 | cellular anatomical entity morphogenesis | + | GO:0098773 | skin epidermis development | - |
| GO:0000902 | cell morphogenesis | + | GO:0031100 | animal organ regeneration | + |
| GO:0048468 | cell development | + | GO:0030154 | cell differentiation | + |
| GO:0048869 | cellular developmental process | + | GO:1990138 | neuron projection extension | + |
| - | - | | GO:0048813 | dendrite morphogenesis | + |
| - | - | | GO:0031016 | pancreas development | + |
| - | - | | GO:0048762 | mesenchymal cell differentiation | + |
| - | - | | GO:0045165 | cell fate commitment | + |
| - | - | | GO:0022612 | gland morphogenesis | + |

| | | | | | |
|---|---|---|---|---|---|
| - | - | | GO:0031175 | neuron projection development | + |
| - | - | | GO:0030097 | hemopoiesis | + |
| - | - | | GO:0007409 | axonogenesis | + |
| - | - | | GO:0030216 | keratinocyte differentiation | + |
| **Custer: Biological process** | | | | | |
| GO:0007155 | cell adhesion | + | GO:0032963 | collagen metabolic process | + |
| GO:0032502 | developmental process | + | GO:0040011 | locomotion | + |
| - | - | | GO:0043473 | pigmentation | + |
| - | - | | GO:0016999 | antibiotic metabolic process | + |
| **Custer: Multicellular organismal process** | | | | | |
| GO:0060249 | anatomical structure homeostasis | + | GO:0022600 | digestive system process | + |
| - | - | | GO:0007631 | feeding behavior | + |
| - | - | | GO:0007612 | learning | + |
| - | - | | GO:0007565 | female pregnancy | + |
| - | - | | GO:0001894 | tissue homeostasis | + |
| - | - | | GO:0050954 | sensory perception of mechanical stimulus | + |
| - | - | | GO:0060135 | maternal process involved in female pregnancy | + |
| - | - | | GO:0007610 | behavior | + |
| - | - | | GO:0007611 | learning or memory | + |
| - | - | | GO:0003014 | renal system process | - |
| - | - | | GO:0007565 | female pregnancy | - |
| - | - | | GO:0048871 | multicellular organismal-level homeostasis | - |
| **Custer: Multicellular organismal process** | | | | | |
| - | - | | GO:0001942 | hair follicle development | + |
| - | - | | GO:0048565 | digestive tract development | + |
| - | - | | GO:0001889 | liver development | + |
| - | - | | GO:0001764 | neuron migration | + |

| | | | | | |
|---|---|---|---|---|---|
| - | - | | GO:0009952 | anterior/posterior pattern specification | + |
| - | - | | GO:0048568 | embryonic organ development | + |
| - | - | | GO:0001525 | angiogenesis | + |
| - | - | | GO:0048706 | embryonic skeletal system development | + |
| - | - | | GO:0001843 | neural tube closure | + |
| - | - | | GO:0021675 | nerve development | + |
| - | - | | GO:0001701 | in utero embryonic development | + |
| - | - | | GO:0007275 | multicellular organism development | + |
| - | - | | GO:0001655 | urogenital system development | + |
| - | - | | GO:0030900 | forebrain development | + |
| - | - | | GO:0048534 | hematopoietic or lymphoid organ development | + |
| - | - | | GO:0030324 | lung development | + |
| - | - | | GO:0007369 | gastrulation | + |
| - | - | | GO:0042063 | gliogenesis | + |
| - | - | | GO:0001822 | kidney development | + |
| - | - | | GO:0048562 | embryonic organ morphogenesis | + |
| - | - | | GO:0030218 | erythrocyte differentiation | + |
| **Unclustered** | | | | | |
| GO:0042098 | T cell proliferation | + | GO:0065008 | regulation of biological quality | + |
| GO:0002694 | regulation of leukocyte activation | + | GO:0014812 | muscle cell migration | + |
| GO:0051480 | regulation of cytosolic calcium ion concentration | + | GO:0001666 | response to hypoxia | + |
| GO:0070663 | regulation of leukocyte proliferation | + | GO:0032879 | regulation of localization | + |
| GO:0032611 | interleukin-1 beta production | + | GO:0007411 | axon guidance | + |
| GO:0005975 | carbohydrate metabolic process | + | GO:0097164 | ammonium ion metabolic process | + |
| GO:0007267 | cell-cell signaling | + | GO:0043406 | positive regulation of MAP kinase activity | + |
| - | - | | GO:0071260 | cellular response to mechanical stimulus | + |

| | | | | | |
|---|---|---|---|---|---|
| - | - | | GO:0000077 | DNA damage checkpoint signaling | + |
| - | - | | GO:0042596 | fear response | + |
| - | - | | GO:0006302 | double-strand break repair | + |
| - | - | | GO:0044774 | mitotic DNA integrity checkpoint signaling | + |
| - | - | | GO:0010522 | regulation of calcium ion transport into cytosol | + |
| - | - | | GO:0035264 | multicellular organism growth | + |
| - | - | | GO:0050851 | antigen receptor-mediated signaling pathway | + |
| - | - | | GO:0006281 | DNA repair | + |
| - | - | | GO:0007166 | cell surface receptor signaling pathway | + |
| - | - | | GO:0051241 | negative regulation of multicellular organismal process | + |
| - | - | | GO:0030317 | flagellated sperm motility | + |
| - | - | | GO:0007187 | G protein-coupled receptor signaling pathway, coupled to cyclic nucleotide second messenger | + |
| - | - | | GO:0044092 | negative regulation of molecular function | + |
| - | - | | GO:0006874 | intracellular calcium ion homeostasis | + |
| - | - | | GO:0019933 | cAMP-mediated signaling | + |
| - | - | | GO:0007189 | adenylate cyclase-activating G protein-coupled receptor signaling pathway | + |
| - | - | | GO:0043312 | neutrophil degranulation | + |
| - | - | | GO:0043603 | amide metabolic process | + |
| - | - | | GO:0050896 | response to stimulus | + |

**Table S4.** Significantly enriched GO terms (*p*-value < 0.05) identified through gene family analyses in the Tropical Eastern Pacific geminate of the damselfish *Azurina atrilobata* and grouper *Cephalopholis colonus*. Enriched expanded and contracted terms are listed by cluster as identified through semantic similarity. Cluster labels assigned to each cluster represent the closest common GO ancestor for that cluster. Expanded enrichment terms are shown with a (+) and contracted a (-). Shared GO terms between the damselfish and groupers are shown in bold.

| Damselfish TEP geminate: *A. atrilobata* | | | Grouper TEP geminate: *C. colonus* | | |
|---|---|---|---|---|---|
| **Custer: Response to chemical** | | | | | |
| - | - | | GO:0010243 | response to organonitrogen compound | - |
| - | - | | GO:0010038 | response to metal ion | - |
| - | - | | GO:0014070 | response to organic cyclic compound | - |
| - | - | | GO:1901700 | response to oxygen-containing compound | - |
| - | - | | GO:0009410 | response to xenobiotic stimulus | - |
| **Custer: Signal transduction** | | | | | |
| GO:0035556 | intracellular signal transduction | - | GO:0007215 | glutamate receptor signaling pathway | - |
| GO:0007166 | cell surface receptor signaling pathway | - | GO:0016055 | Wnt signaling pathway | - |
| GO:0006954 | inflammatory response | + | - | - | |
| GO:0009605 | response to external stimulus | + | - | - | |
| **Custer: Primary metabolic process** | | | | | |
| GO:0034654 | nucleobase-containing compound biosynthetic process | - | GO:0006470 | protein dephosphorylation | - |
| GO:0006163 | purine nucleotide metabolic process | - | - | - | |
| GO:1901566 | organonitrogen compound biosynthetic process | + | - | - | |
| GO:0006508 | proteolysis | + | - | - | |
| GO:0044271 | cellular nitrogen compound biosynthetic process | + | - | - | |
| GO:0018193 | peptidyl-amino acid modification | + | - | - | |
| **Custer: Regulation of metabolic process** | | | | | |

| | | | | | |
|---|---|---|---|---|---|
| GO:0006355 | regulation of DNA-templated transcription | + | GO:0031323 | regulation of cellular metabolic process | - |
| - | - | | GO:0080090 | regulation of primary metabolic process | - |
| **Custer: Biological regulation** | | | | | |
| GO:0065008 | regulation of biological quality | + | GO:0010975 | regulation of neuron projection development | - |
| GO:0042127 | regulation of cell population proliferation | + | - | - | |
| **Custer: Regulation of cell differentiation** | | | | | |
| **GO:0050767** | **regulation of neurogenesis** | + | **GO:0050767** | **regulation of neurogenesis** | - |
| - | - | | GO:0022604 | regulation of cell morphogenesis | - |
| - | - | | GO:0044057 | regulation of system process | - |
| - | - | | GO:0045664 | regulation of neuron differentiation | - |
| **Custer: Positive regulation of cell differentiation** | | | | | |
| GO:0010720 | positive regulation of cell development | + | GO:0045666 | positive regulation of neuron differentiation | - |
| GO:0051962 | positive regulation of nervous system development | + | GO:0050769 | positive regulation of neurogenesis | - |
| **Custer: Negative regulation of metabolic process** | | | | | |
| GO:0031327 | negative regulation of cellular biosynthetic process | - | - | - | |
| GO:0045934 | negative regulation of nucleobase-containing compound metabolic process | - | - | - | |
| GO:0045936 | negative regulation of phosphate metabolic process | - | - | - | |
| GO:0031400 | negative regulation of protein modification process | + | - | - | |
| GO:0045936 | negative regulation of phosphate metabolic process | + | - | - | |
| GO:0010629 | negative regulation of gene expression | + | - | - | |
| **Custer: Transport** | | | | | |
| GO:0006810 | transport | + | GO:0051668 | localization within membrane | - |
| GO:0015833 | peptide transport | + | GO:0098655 | monoatomic cation transmembrane transport | - |
| GO:0033365 | protein localization to organelle | + | GO:0006887 | exocytosis | - |

| | | | | | |
|---|---|---|---|---|---|
| GO:0006886 | intracellular protein transport | + | GO:0051656 | establishment of organelle localization | - |
| **Custer: Regulation of transport** | | | | | |
| GO:0002791 | regulation of peptide secretion | + | GO:0010959 | regulation of metal ion transport | - |
| - | - | | GO:0060341 | regulation of cellular localization | - |
| **Custer: Cellular component organization** | | | | | |
| GO:0034330 | cell junction organization | + | GO:0065003 | protein-containing complex assembly | - |
| GO:0006996 | organelle organization | + | GO:0050808 | synapse organization | - |
| - | - | | GO:0061024 | membrane organization | - |
| - | - | | GO:0051276 | chromosome organization | - |
| **Custer: Anatomical structure development** | | | | | |
| **GO:0060429** | **epithelium development** | + | **GO:0060429** | **epithelium development** | + |
| **GO:0009887** | **animal organ morphogenesis** | + | **GO:0009887** | **animal organ morphogenesis** | + |
| GO:0048646 | anatomical structure formation involved in morphogenesis | + | GO:0061061 | muscle structure development | + |
| GO:0048589 | developmental growth | + | GO:0048812 | neuron projection morphogenesis | - |
| **Custer: Developmental process** | | | | | |
| **GO:0048731** | **system development** | + | **GO:0048731** | **system development** | + |
| GO:0001654 | eye development | + | GO:0007275 | multicellular organism development | + |
| GO:0048666 | neuron development | + | GO:0048667 | cell morphogenesis involved in neuron differentiation | - |
| **Unclustered** | | | | | |
| GO:1902532 | negative regulation of intracellular signal transduction | + | GO:0044703 | multi-organism reproductive process | + |
| GO:0043603 | amide metabolic process | + | GO:0007283 | spermatogenesis | + |
| GO:0044092 | negative regulation of molecular function | + | GO:0032501 | multicellular organismal process | + |
| GO:0051050 | positive regulation of transport | + | GO:0022414 | reproductive process | + |
| GO:0040011 | locomotion | + | GO:0044772 | mitotic cell cycle phase transition | - |
| GO:0071363 | cellular response to growth factor stimulus | + | GO:0048167 | regulation of synaptic plasticity | - |
| GO:0050900 | leukocyte migration | + | GO:0050905 | neuromuscular process | - |

| | | | | | |
|---|---|---|---|---|---|
| GO:0070372 | regulation of ERK1 and ERK2 cascade | + | GO:0050803 | regulation of synapse structure or activity | - |
| GO:0048871 | multicellular organismal-level homeostasis | + | GO:0071214 | cellular response to abiotic stimulus | - |
| GO:0043066 | negative regulation of apoptotic process | + | GO:0060079 | excitatory postsynaptic potential | - |
| GO:1902533 | positive regulation of intracellular signal transduction | + | GO:0042981 | regulation of apoptotic process | - |
| - | - | | GO:0030003 | intracellular monoatomic cation homeostasis | - |
| - | - | | GO:0009314 | response to radiation | - |
| - | - | | GO:0032535 | regulation of cellular component size | - |
| - | - | | GO:0010564 | regulation of cell cycle process | - |
| - | - | | GO:0010976 | positive regulation of neuron projection development | - |
| - | - | | GO:0009057 | macromolecule catabolic process | - |
| - | - | | GO:0098609 | cell-cell adhesion | - |
| - | - | | GO:0043085 | positive regulation of catalytic activity | - |

**Table S5.** Additional damselfish and grouper species used in positive selection analyses and gene family expansion-contraction analyses. Genbank accession numbers given for the genome assemblies used and the RNAseq data used for evidence in annotation of the genomes.

| **Species** | **Assembly** | **RNAseq** |
|---|---|---|
| *Acanthochromis polyacanthus* | GCF_021347895.1 | SRR15693664, SRR15693670, SRR15693672, SRR15693676, SRR15693678, SRR15693680, SRR15693682, SRR15693684, SRR15693689, SRR15693691 |
| *Amphiprion clarkii* | GCA_027123335.1 | ERR2704777, ERR2704778, ERR2704779, ERR2704780, ERR2704781, ERR2704782, ERR2704783, ERR2704784, ERR2704785, ERR2704786, ERR2704787, ERR2704788, ERR2704789, ERR2704790, ERR2704791, ERR2704792, ERR2704793, ERR2704794 |
| *Amphiprion ocellaris* | GCF_022539595.1 | SRR23998317, SRR23998318, SRR23998319, SRR23998320, SRR23998321, SRR23998322, SRR23998323, SRR23998324, SRR23998325, SRR23998326, SRR23998327, SRR23998328, SRR23998329, SRR23998330, SRR23998331, SRR23998332, SRR23998333, SRR23998334, SRR23998335, SRR23998336 |

| | | |
|---|---|---|
| *Amphiprion percula* | GCA_003047355.2 | ERR2704783, ERR2704784, ERR2704785, ERR2704786, ERR2704788 |
| *Dascyllus trimaculatus* | GCA_024666655.1 | SRR5997703 |
| *Abudefduf troschelii* | *See Tracy et al. 2025* | |
| *Abudefduf saxatilis* | *See Tracy et al. 2025* | |
| *Centropristis striata* | GCF_030273125.1 | SRR13271205 |
| *Cromileptes altivelis* | GCA_013133815.1 | SRR17595877 |
| *Epinephelus cyanopodus* | GCA_026686955.1 | - |
| *Epinephelus fuscoguttatus* | GCF_011397635.1 | SRR19852718, SRR19852719, SRR19852720, SRR19852721, SRR19852722, SRR19852723 |
| *Epinephelus lanceolatus* | GCF_005281545.1 | SRR17978194, SRR17978195, SRR17978196, SRR17978197, SRR17978198, SRR17978199, SRR17978200, SRR17978201, SRR17978202, SRR17978203, SRR17978204, SRR17978205 |
| *Epinephelus moara* | GCF_006386435.1 | SRR9886795, SRR9886796, SRR9886797, SRR9886798, SRR9886799, SRR9886800, SRR9886801, SRR9886802, SRR9886803 |
| *Plectropomus leopardus* | GCF_008729295.1 | SRR16351828, SRR16351829, SRR16351830, SRR16351832, SRR16351833, SRR17640161, SRR17640162, SRR17640163, SRR17640164, SRR17640165, |

| | | |
|---|---|---|
| | | SRR17640166, SRR18762158, SRR19285154, SRR19285155, SRR19285156, SRR19285157, SRR19285158, SRR19285159, SRR19660399, SRR19660400, SRR19660401, SRR19660402, SRR19660403, SRR19660404, SRR19660406, SRR19660407, SRR19660408, SRR19660409 |

**Table S6.** log-likelihood scores and lambda estimates for the multiple values of *gamma* rate categories trialled in CAFE5 analysis for the damselfish and groupers. Increasing values for *gamma* were attempted until the log-likelihood score showed no improvement.

| Family | *gamma* categories: | Final Likelihood (-lnL) | Lambda |
|---|---|---|---|
| *Damselfish* | base | 172963 | 0.0031883 |
| *Damselfish* | 2 | 162317 | 0.0088295 |
| *Damselfish* | 3 | 160605 | 0.0043469 |
| *Damselfish* | 4 | 159076 | 0.0053617 |
| *Damselfish* | 5 | 159868 | 0.0037713 |
| *Groupers* | base | 147406 | 0.0013059 |
| *Groupers* | 2 | 139530 | 0.0036187 |
| *Groupers* | 3 | 138153 | 0.0026773 |
| *Groupers* | 4 | 142288 | 0.0007854 |

| | | | |
|---|---|---|---|
| *Groupers* | 5 | 138041 | 0.0027123 |
| *Groupers* | 6 | 140052 | 0.0011307 |

**Table S7.** Convergence of ten repeat runs of CAFE5 analyses in the damselfish at *gamma* = 2.

| Run | Likelihood (-lnL): | Lambda: | Epsilon: | Max lambda: | Alpha: |
|---|---|---|---|---|---|
| 1 | 162317.01668889 | 0.008828 | 0.036057 | 0.022001 | 0.401085 |
| 2 | 162316.40000330 | 0.008828 | 0.036057 | 0.022001 | 0.401095 |
| 3 | 162316.99741819 | 0.008828 | 0.036057 | 0.022001 | 0.401093 |
| 4 | 162316.99398100 | 0.008828 | 0.036057 | 0.022001 | 0.401094 |
| 5 | 162317.01451515 | 0.008829 | 0.036057 | 0.022001 | 0.401065 |
| 6 | 162317.01544645 | 0.008829 | 0.036057 | 0.022001 | 0.401082 |
| 7 | 162316.99631608 | 0.008828 | 0.036057 | 0.022001 | 0.401091 |
| 8 | 162317.01419810 | 0.008828 | 0.036057 | 0.022001 | 0.401096 |
| 9 | 162317.00673940 | 0.008827 | 0.036057 | 0.022001 | 0.401087 |
| 10 | 162317.00991234 | 0.008828 | 0.036057 | 0.022001 | 0.401103 |

**Table S8.** Convergence of ten repeat runs of CAFE5 analyses in the groupers at *gamma* = 2.

| Run | Likelihood (-lnL): | Lambda: | Epsilon: | Max lambda: | Alpha: |
|---|---|---|---|---|---|
| 1 | 139316.30712603 | 0.004026 | 0.0626 | 0.0125 | 0.2165 |
| 2 | 139316.31944754 | 0.004026 | 0.0626 | 0.0125 | 0.2165 |

| | | | | | |
|---|---|---|---|---|---|
| 3 | 139316.32560927 | 0.004026 | 0.0626 | 0.0125 | 0.2165 |
| 4 | 139316.32048987 | 0.004026 | 0.0626 | 0.0125 | 0.2165 |
| 5 | 139316.31496698 | 0.004026 | 0.0626 | 0.0125 | 0.2165 |
| 6 | 139316.32125573 | 0.004026 | 0.0626 | 0.0125 | 0.2165 |
| 7 | 139316.32742688 | 0.004026 | 0.0626 | 0.0125 | 0.2165 |
| 8 | 139316.30560268 | 0.004026 | 0.0626 | 0.0125 | 0.2165 |
| 9 | 139316.29848339 | 0.004027 | 0.0626 | 0.0125 | 0.2165 |
| 10 | 139316.32036870 | 0.004026 | 0.0626 | 0.0125 | 0.2165 |
